# An anti-amyloidogenic approach to specifically block memory consolidation in mice for therapeutic intervention

**DOI:** 10.1101/2025.01.27.635080

**Authors:** Paula López-García, Kerry R. McGreevy, Maria Eugenia Vaquero, Daniel Ramírez de Mingo, Anna Pallé, Helena Akiko Popiel, Andrea Santi, Yoshitaka Nagai, José Luis Trejo, Mariano Carrión-Vázquez

**Affiliations:** Instituto Cajal (IC-CSIC), Department of Molecular, Cellular and Developmental Neurobiology. IC-CSIC, Avenida Doctor Arce 37, 28002 Madrid, Spain; Instituto Cajal (IC-CSIC), Department of Traslational Neuroscience. IC-CSIC, Avenida Doctor Arce 37, 28002 Madrid, Spain; National Center of Neurology and Psychiatry. National Institute of Neuroscience, Department of Degenerative Neurological Diseases. 4-1-1 Ogawa-Higashi, Kodaira, Tokyo 187-8502, Japan; Johns Hopkins University, Department of Psychological & Brain Sciences 3400 N. Charles Street, Baltimore, MD 21218, Maryland, USA; Osaka University. Graduate School of Medicine. Department of Neurotherapeutics. 2-2 Yamadaoka, Suita, Osaka 565-0871, Japan

## Abstract

Post-traumatic stress disorder (PTSD) is a mental health disorder triggered by the exposure to a traumatic event, which manifests with anguish, intrusive memories, and negative mood changes. So far, there is no efficient treatment for PTSD other than symptomatic palliative care. However, the cytoplasmic polyadenylation element binding protein-3 (CPEB3) has been recently associated to a potential risk gene for PTSD. Considering that CPEB3 protein is a functional amyloid whose importance for long-term memory consolidation in mammals is well established, we propose the active (amyloid) state of CPEB3 as a promising therapeutic target to block the consolidation of traumatic memories through by the anti-amyloidogenic polyglutamine binding peptide 1 (QBP1).

Here we report a preclinical development in mice of this pharmacological treatment for PTSD based on the action of the QBP1 peptide. We first characterized the human CPEB3 (hCPEB3) protein *in vitro*, showing how its amyloid aggregation is inhibited by the active core of QBP1 (QBP1-M8) without affecting other self-assembly processes such as phase separation. Then, we generated and characterized a novel transgenic mouse that constitutively expresses QBP1 in tandem (TgQBP1). TgQBP1 mice have shown that the consolidation of simple learning is impaired after 24 h for both hippocampal-dependent and aversive memories and that it is limited to new learned memories and has no effect on short-term memory. Furthermore, fear induced anxiety was reduced in comparison to WT mice, suggesting that PTSD-like symptoms are also being ameliorated. Intriguingly, we found that aversive memories seem to be more strongly affected in younger mice. Finally, the analysis of CPEB3 amyloid presence in hippocampal extracted samples showed a correlative decrease in murine CPEB3 oligomerization in the TgQBP1 mice. Taking together, these results strongly suggest that the amyloidogenic blockage of CPEB3 by QBP1 peptide is a potential drug for the prevention and treatment of PTSD.

## Introduction

<u>P</u>ost-traumatic <u>s</u>tress <u>d</u>isorder (PTSD) is a mental disorder classified within the group of disorders related to stress factors ^1^, in which exposure to a traumatic event arises specific symptoms (intrusive memories, impulsive behaviors, hyperexcitation, anxiety and mood changes) that result in a serious alteration of the patient’s life ^2–5^. Within the first days and weeks after the trauma, <u>a</u>cute <u>s</u>tress <u>d</u>isorder (ASD) can develop as a first disturbance of the stress response ^6–8^, but in an unfortunate 5-8% of the general population (which is increased in war veterans, firefighters and related jobs) these symptoms persist for more than 6 months and finally develop PTSD ^9–11^. In terms of gender, the impact ratio is 1:20 in men and 1:10 in women. The factors that influence the probability of suffering this disorder are the intensity, duration and proximity of exposure to the traumatic event ^12–14^.

During the last four decades, most of the pharmacology for this disorder was derived from drugs already on the market for diseases such as anxiety disorders and depression ^3,15,16^. Only selective serotonin reuptake inhibitors (SSRIs), such as Sertraline, Paroxetine and Fluoxetine, demonstrated efficacy in the treatment of PTSD ^17,18^. However, they failed to achieve complete recovery of patients and were mainly applied to alleviate symptoms in combination with psychotherapy ^14,19^. Additionally, the use of other antidepressants such as Mirtazapine and Venlafaxine was introduced, which showed similar efficacy to SSRIs ^17,20,21^ although the corresponding clinical trials were limited due to the heterogeneity of PTSD and the complexity of its symptoms ^22^. On the other hand, psychological interventions still remain as the most effective PTSD treatment, specifically <u>c</u>ognitive <u>b</u>ehavioral therapy (CBT) coupled with focused-trauma approaches ^16,23,24^. Related to this, in recent years memory itself has been raised as a therapeutic target towards a specific treatment for PTSD ^25–28^, since the primary cause of the disorder lies in the prolonged stress caused by a traumatic event and how the recall of this memory is encoded as a context (location, smell, sound…) associated with a strong negative emotional valence (feelings of pain, fear, anguish) ^28–31^. As a result, once the disorder has been developed, any stimulus from the environment could induce an unconscious recall of traumatic events in a safe context ^3,29,32^, triggering a crisis. Modulating those stimuli and emotional cues during active trauma recall was the main focus of these new therapies ^33–35^, acting specifically on fear conditioning and extinction of traumatic events. Currently, the most promising preventive pharmacology is based on the <u>h</u>ypothalamic–<u>p</u>ituitary–<u>a</u>drenal (HPA) axis and *β*-adrenergic blockers ^16,36^; thus, hydrocortisone ^25,37,38^, oxytocin ^39–41^ and propranolol ^42–45^ are able to reduce the impact of fear memories (in human and rodents) and ameliorate PTSD symptoms, but they are still more focused on reconsolidation-based approaches in combination with psychological interventions ^23,25,42,44,46^ than being a first-line solution itself.

Based on this approach, one could expect that directly interfering on protein synthesis and synaptic plasticity would be a more effective strategy, as it will directly affect the very first consolidation process after the trauma exposure. In the search for new specific therapeutic targets related to memory consolidation and PTSD onset, the CPEB family stands out as the most promising one since they are RNA-binding proteins cable of regulating mRNA translation and mediate memory storage and synaptic plasticity ^47–49^, along which CPEB3 isoform is considered as a potential risk gene for PTSD ^50^. In vertebrates, CPEB is usually coded by four genes of which the first member identified was CPEB1, with CPEBs 2-4 later described in mice ^51–54^. This family of proteins share a RNA-binding domain (RBD) at their C-terminal regions and in the case of CPEB2 and CPEB3 there are N-terminal regions rich in glutamine/asparagine residues (Q/N regions), which resemble the amyloid/prionoid Q/N region of their orthologues in *Aplysia* and *Drosophila* (*Ap*CPEB and Orb2, respectively) ^51,55–57^. CPEB3 is able to form self-propagating and heritable aggregates in yeast and its aggregation in mice brain occurs only under physiological conditions after neuronal stimulation, associated with long-term memory (LTM) ^56,58^. Furthermore, CPEB3 <u>k</u>nock <u>o</u>ut mice (KO) showed defects in memory consolidation associated with spatial learning and the interruption in CPEB3 function after consolidation result in an impairment in long-term memory ^58–60^. CPEB2 is also required for long-term memory consolidation ^61,62^, but its amyloidogenic and prionoid properties has not been characterized yet. Considering the functional importance of the Q/N prionoid regions of the CPEB family, their evolutionary conservation ^47,52^ and their function as a switch for long-term memory through protein synthesis in mammals, CPEB3 constitutes a promising therapeutic target for pharmacological interventions for PTSD.

Polyglutamine binding peptide 1 (QBP1: SNWKWWPGIFD) is an anti-amyloidogenic dodecapeptide originally developed to specifically bind to long polyQ stretches (*i.e.,* pathological) ^63,64^, as a therapeutic approach for neurodegenerative diseases known as polyglutaminopathies ^65,66^. According to the proposed mechanism of action, QBP1 specifically binds to the protein monomer and is cable of blocking the transition of the toxic β-sheet rich conformation that initiates amyloidogenesis ^67,68^, thus blocking this process at its start. In addition to polyQ regions, QBP1 has shown polyvalence by blocking the conformational polymorphism of polyQ, Sup35NM, and α-synuclein at the monomer level ^67,69^, as well the amyloidogenesis process of TDP-43 ^70^. Interestingly, it was also demonstrated that the minimum active core named QBP1-M8 (octapeptide: WKWWPGIF) ^71,72^ also inhibits *in vitro* the amyloidogenesis of functional amyloids like *Ap*CPEB ^73^ and Orb2 ^74^. Also, a preliminary report on the action of QBP1 on CPEB3 was recently advanced ^75^. Moreover, the expression of a tandem QBP1 construct in a *Drosophila* transgenic model showed that it is able to block memory consolidation and learning ^74^. Based on all this, we proposed that the glutamine based structures present in murine CPEB3 could also be blocked by QBP1, allowing us to develop a specific pharmacological approach for PTSD.

Here, we have first characterized the interplay between QBP1 and CPEB3 protein *in vitro*, confirming the inhibition of the amyloid state and analyzing the nature of their interaction. Second, we have generated and characterized a transgenic mouse constitutively expressing QBP1 in tandem (TgQBP1 mice), showing an impairment in long-term memory accompanied by a decreased mCPEB3 aggregation at hippocampal samples in the presence of QBP1. Our results provide a proof of concept of a novel and specific pharmacological approach to block memory consolidation of traumatic memories in mammals by QBP1, which could serve for preventing and treating PTSD and ASD.

## Results

### Comprehensive characterization of QBP1 inhibition of human CPEB3 amyloidogenesis

The effect of QBP1 on human CPEB3 amyloidogenesis was recently advanced in our previous work ^75^, where QBP1 was mostly used as an amyloid modulator. Here we have fully characterized its inhibitory effects on amyloidogenesis. We first examined the amyloid kinetics of the hCPEB3 intrinsically disorder region (hIDR_1-450_)^75^ by Thioflavin T (ThT) fluorescence intensity in the presence of either the QBP1 active core (QBP1-M8, WKWWPGIF) or its *Scrambled* version (SCR-C: WPIWKGWF) as a negative control ^64,71,72^. Amyloid kinetics in the presence of QBP1-M8 and SCR-C at a ratio of 1:8 (protein:inhibitor) showed that both peptides were able to inhibit hIDR fibrillation (**Fig. 1A**). We further analyzed whether QBP1-M8 inhibition affects the different intermediate amyloid species formed by hIDR using specific conformational antibodies against different amyloid species within a immunodot blot and SDD-AGE, performed in presence of slight agitating to minimize unspecific aggregation. The immunodot blot analysis of hIDR incubated with A11/OC conformational antibodies^76^ showed an inhibition of the formation of pre-fibrilar oligomers and mature oligomers and fibers in the samples containing QBP1-M8 (**Fig. 1B**). The SDD-AGE analysis showed also a reduction in the formation of aggregates/oligomers smears in the presence of QBP1-M8 (**Fig. 1C**). The species formed throughout the aggregation cascade were then visualized in presence of QBP1-M8. In the TEM micrographs of hIDR we observed the formation of non-specific aggregates mixed with some fibrils even at 216h when QBP1-M8 is present (**Fig. 1D**), with a reduction of fiber formation indicative of the inhibition of its amyloidogenesis. In the case of the amiloidogenic core region ^75^, named hCPEB3 R1 (residues 1-200, **Fig. S1A**), we observed enhanced non-specific aggregation and fibril-like assemblies still present, which is consistent with an enhancement of its aggregation kinetics in the presence of QBP1-M8 (**Fig. 1E**). Moreover, a longer amyloid region of the hCPEB3 IDR (named R2 ^75^, spanning 1-300 residues) showed abundant fibrillation *in vitro,* which in the presence of QBP1-M8 showed a large amount of protofibrils evolving into amyloid fibers mixed with non-specific aggregates (**Fig. S1B)**.

**Figure 1.**
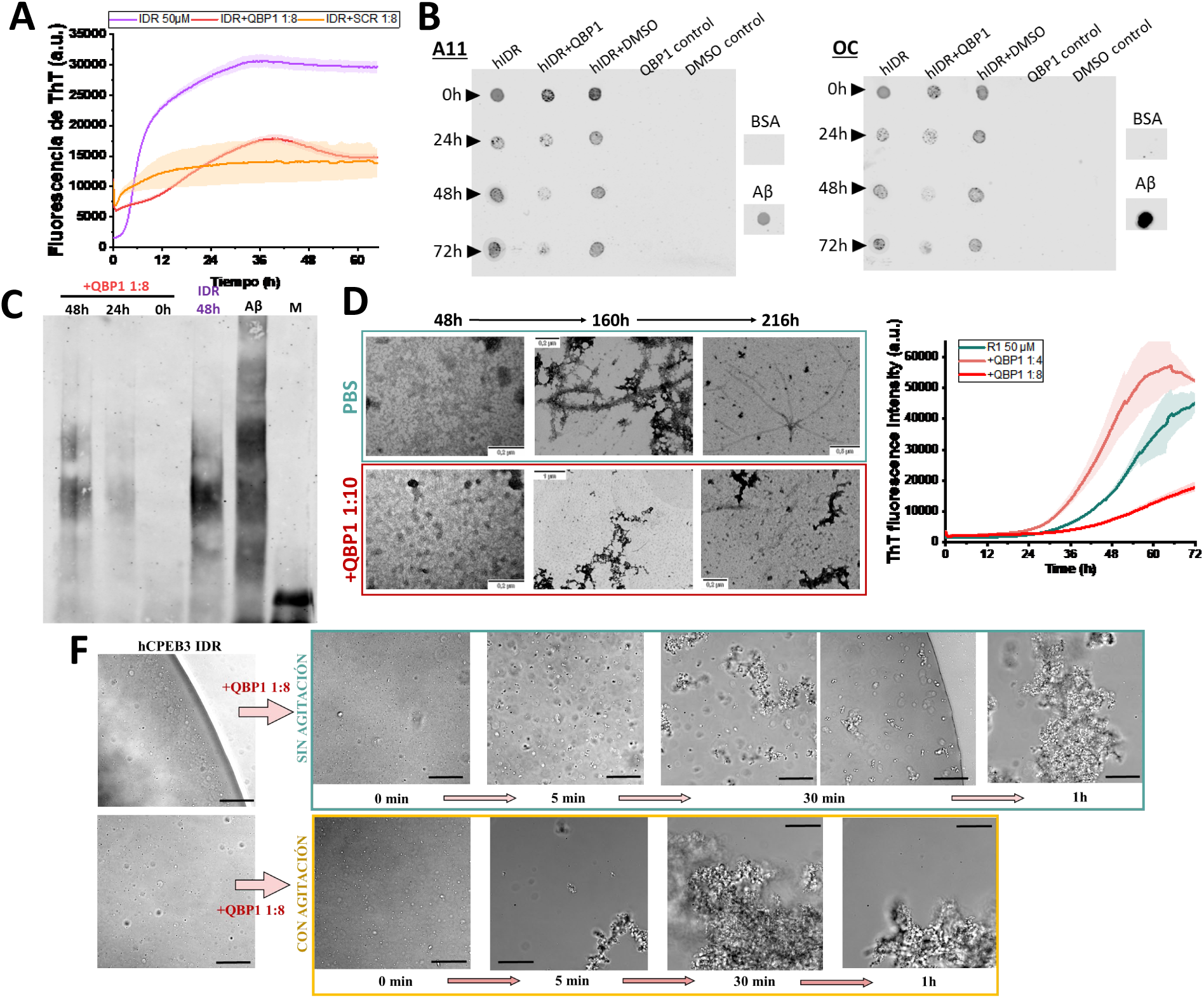
QBP1-M8 inhibits the amyloidogenesis of hCPEB3 IDR and its fragments. A) Kinetics of amyloid aggregation of hCPEB3 IDR (50 μM) monitored by ThT fluorescence emission in the presence or absence of QBP1-M8 (1:8 ratio) and SCR-C (1:8 ratio). Error bars indicate the standard error of the mean (n=3) of a representative experiment. B) Immunodot blot of hCPEB3 IDR (50 μM, purple) in the presence or absence of QBP1-M8 (1:8, red) or DMSO (vehicle, 1:8, yellow) which reflects the presence of A11/OC reactive species formed after incubation at 37 ᵒC with shaking (800 rpm thermoblock). Aβ and BSA were used as positive and negative controls, respectively. C) SDD-AGE of hCPEB3 IDR (40 μM) in the presence or absence of QBP1 (1:8) at different time points of incubation at 37 ᵒC with shaking (800 rpm, thermoblock) and incubated with an anti-OC antibody (against fibrillar oligomers and fibers). Aβ was used as a positive control. D) Representative images of hIDR (10 μM) fibrillation in the presence or absence of QBP1-M8 (1:10). Samples were incubated at 37 ᵒC in PBS pH 7.4 without shaking. The image scale in each sample is indicated. E) Kinetics of hCPEB3 R1_1-200_ (50 μM) amyloid aggregation, monitored by ThT fluorescence emission, in the presence or absence of QBP1-M8 (1:4-1:8). Error bars indicate the standard error of the mean (n = 3) of a representative experiment. F) Phase-contrast micrographs of the effect of QBP1-M8 (1:8 ratio) on the phase separation of hCPEB3 IDR 50 μM upon co-incubation in the absence (upper turquoise panel) or presence of agitation (37°C and 800 rpm, lower yellow panel) under physiological-like conditions (PBS pH 7.4). Scale bar: 0.5 μm.

Finally, we examined whether QBP1-M8 peptide affects the phase separation behavior of hCPEB3 IDR ^75^. The addition of QBP1-M8 (1:8 ratio) to liquid droplets of this protein incubated without agitation showed that phase separation is maintained over time until formation of aggregates saturates the protein concentration (**Fig. 1F**, upper turquoise panel). Upon agitated co-incubation, nonspecific aggregates formed earlier (**Fig. 1F**, lower yellow panel). These results suggest that QBP1-M8 lowers amyloid aggregation without altering the phase separation process.

### QBP1 transgenic mice showed impaired long-term memory

Once we demonstrated *in chemico* that QBP1-M8 inhibits the active amyloid state of CPEB3, we set out to test the *in vivo* proof of concept that QBP1 could maintain its inhibitory effect and in turn block memory consolidation in a mammalian model. To that end, we generated a transgenic mouse constitutively expressing QBP1 in tandem (SNWKWWPGIFD-SNWKWWPGIFD), to further increase its activity^68,77^, fused to the <u>g</u>reen fluorescence <u>p</u>rotein (eGFP) (TgQBP1, **Fig. S2A-B)**. After verifying the presence of QBP1 in the hippocampus of the TgQBP1 mice **(Fig. S2C)**, we performed an extensive battery of general health and developmental tests in order to discard gross defects **(Fig. S2D)**.

First, we characterized the locomotor activity of TgQBP1 mice by analyzing them in a two-day protocol (**Fig. 2A**): while Day 1 measures the locomotor activity itself, where the animal freely explores the apparatus cage, its re-exposure on Day 2 (24 h later) reflects their exploratory behavior in a familiar environment (usually lower). The horizontal activity showed normal locomotor activity with no differences compared to control mice (WT) on Day 1 (**Fig. 2B**), although TgQBP1 mice were significantly more active than WT mice on Day 2 (p<0.001), even if both groups significantly decreased their activity compared to the previous day (WT p<0.001 and TgQBP1 p=0.017). This behavior was similar for the vertical activity (**Fig**. **S3A**), where TgQBP1 mice significantly displayed an increased activity on Day 2 (p=0.027). Indeed, the horizontal activity by minute evinced that mice varied their activity according to the minute, day and group (p=0.028; **Fig. 2C**). However, on Day 1 mice decreased the activity over time regardless of group (p<0.001), but on Day 2 TgQBP1 mice had significantly higher overall activity than WT mice (p<0.001) and specifically in the last minutes of the test (Min3 p<0.001; Min4 p=0.003; Min5 p=0.01). Then, to examine whether this effect relied on different basal levels of anxiety, we performed an Elevated plus maze test. We did not found significant differences in any of the analyzed variables (**Fig. S3B-C**), which suggests normal levels of anxiety in the TgQBP1 mice. Taken together, we propose that abnormal re-exploratory behavior of the TgQBP1 mice could be originated from an impairment in the memory consolidation of the known environment.

**Figure 2.**
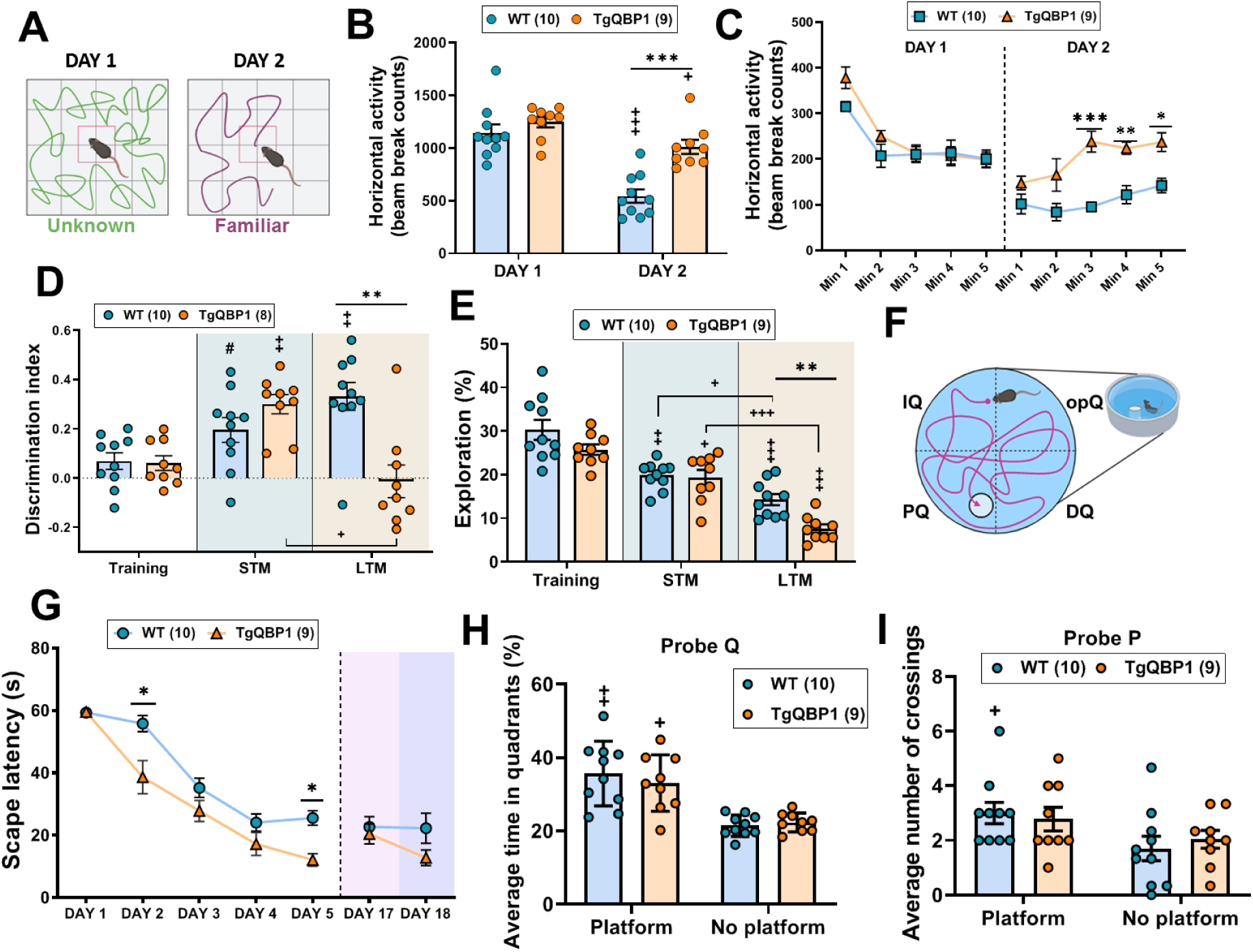
Hippocampal-dependent memory is impaired in the TgQBP1 mice. **A)** Representative schematic of the two-day activity cage protocol: the first day, the locomotor activity is assessed in a novel environment, while in the second day the locomotor activity is analyzed in a familiar environment. **B)** Total horizontal activity on both days of the protocol in the activity cage. On Day 1, no differences between groups were observed, while re-exposure on Day 2 showed that TgQBP1 mice significantly increased their activity compared to WT mice (p<0.001). Furthermore, both groups significantly decreased their activity compared to Day 1 (WT p<0.001 and TgQBP1 p=0.017). **C)** The horizontal activity per minute on each test day reflected that mice modify their behavior as a function of day, minute and group (Interaction p=0.028). On Day 1, there were no significant differences between groups, although both groups changed their behavior significantly over minutes (Time p<0.001). On Day 2, there was no interaction, but both groups varied their behavior over time (p=0.0014) and as a function of group (p<0.001). Specifically, TgQBP1 mice showed significantly more activity than WT mice in the last minutes of the test (p<0.001; p=0.003 and p=0.01 respectively). **D)** Discrimination Index (DI) values are shown for each phase of the test. WT mice tended to show higher DI scores in the STM phase (p=0.059) and significantly higher DI scores in the Long-Term Monitoring (LTM) phase (p=0.007). TgQBP1 mice showed significantly higher DI scores in the Short-term Memory (STM) phase compared to the training phase (p=0.01), but showed significantly lower DI scores in the LTM phase (p=0.025) and lowered significantly their DI scores in the LTM phase compared to the STM phase (p=0.012). Finally, TgQBP1 mice tended to show higher DI scores than WT animals in the STM phase (p=0.057) and significantly lower DI scores in the LTM phase (p=0.002). **E)** The exploration percentage reflects that there is a significant interaction between the time of the test and the group (p=0.049). WT mice significantly decreased their exploration with respect to training (STM p=0.002 and LTM p<0.001), as well as significantly decreased from short to long-term (p=0.018). TgQBP1 mice also significantly decreased their exploration compared to training (STM p=0.016 and LTM p<0.001) and between short and long-term (p<0.001). However, it stands out that TgQBP1 mice explored significantly less the novel object in the long-term (p=0.002). **F)** Representative diagram of the Morris Water Maze. This test consists of evaluating the spatial and cognitive memory of animals, which must find a transparent platform located in one of the four quadrants of a pool (PQ). For several days, four times each day, the mice learn the location of the platform and take increasingly less time to leave (escape latency). **G)** Acquisition of platform localization, where escape latency decreased over days in both groups, especially in the TgQBP1 mice (Day 2 p<0.001 and Day 5 p<0.001). However, LTM at 17-18 days reflected a loss of that improvement (pink-lilac). **H)** Percentage of time in each quadrant after acquisition (Probe Q), in which both groups spent significantly more time in PQ than in the average of the other quadrants (WT p=0.004 and TgQBP1 p=0.014). **I)** Mean number of crossings by platform location (Probe P), in which only WT mice significantly crossed more times the exact location of the platform in the PQ quadrant than the TgQBP1 mice with respect to the same location in the other quadrants (p=0.017). For comparisons between independent groups: *p<0.05, **p<0.01, ***p<0.001, trends 0.05 ≥ # < 0.09; for comparisons between dependent groups: +p<0.05, ++p<0.01, +++p<0.001, trends 0.05 ≥ #’ < 0.09

Next, we analyzed the integrity of the hippocampus-dependent memories in TgQBP1 mice. A novel object recognition test was performed to evaluate rodents’ ability to recognize a novel object in the absence of reinforcement (**Fig. S3D**), where they should spend more time exploring the novel object as they have a natural preference for novelty ^78^. According to the discrimination index (DI, exploration of the new object compared to total exploration), both groups equally explored the two unfamiliar objects in the training phase (**Fig. 2D**). One hour later (short term, **Fig. 2D** turquoise), the two groups explored significantly more the novel object according to training (WT p=0.089 and TgQBP1 p=0.004). After 24 h (long-term, **Fig. 2D** beige), WT mice improved their discrimination in comparison to training (p=0.002), while TgQBP1 mice significantly worsened it in comparison to short term (p=0.041), showing a significant difference between the two groups (p=0.003). Overall, TgQBP1 mice were able to discriminate novelty at short term (1h) but their performance was completely deteriorated at long-term (24 h), exploring both objects equally as if they were completely unfamiliar (**Fig. 2D**). In terms of exploration (number of times the animal approaches the object frontally), the percentage reflected that there were significant differences depending on the group and test phase (p=0.049; **Fig. 2E**), and they also needed fewer explorations to discriminate the novel object (p<0.001). However, TgQBP1 mice explored significantly less than WT mice at long-term (p=0.002), suggesting that their lack of discrimination makes them to spend more time within each exploration. Finally, the percentage of novelty preference (**Fig. S3E**), in which a 50% of exploration means an equal interest in both objects (purple dashed line), confirmed that TgQBP1 mice did not significantly discriminate the novelty after 24 h (long-term, p=0.003), confirming the memory impairment at the long-term phase.

### TgQBP1 mice showed an impaired memory consolidation despite repetitive learning

Considering that the function CPEB3 is based on its conversion to an active amyloid state and that its prion-like properties promote temporal stability, we decided to assess memory consolidation in a repetitive and more complex learning paradigm. The Morris water maze is a hippocampal-dependent task widely used to study spatial learning and memory in rodents, in which mice learn to escape from the pool by finding a transparent platform along four trials per day (**Fig. 2F**). In the acquisition phase, both groups significantly decreased their escape latency over days (Days p<0.001), showing that they had learned the task (**Fig. 2G**). Surprisingly, TgQBP1 mice seemed to performed significantly better than WT mice over time (Group p=0.001) and they performed significantly better on the last day of acquisition (Day 5, p<0.001) (**Fig. 2G**), likely because WT mice performance reach a *plateau* on that final day.

After the acquisition, we analyzed the recall of the platform location. Specifically, we analyzed the time spent in each quadrant (Probe Q, from *quadrant*), where both groups spent significantly more time in the quadrant with the platform (PQ, WT p=0,004 y TgQBP1 p=0,014) (**Fig. 2H**). However, when we analyzed the number of times they have crossed the platform location in that quadrant at the millimeter level (Probe P, from *platform*), only WT mice spent significantly more time on the platform than off the platform (p=0.017; **Fig. 2I**). Taken together, these results suggest that TgQBP1 mice remember the quadrant of the platform, but not the position of the platform within the quadrant, which evidences some impairment of memory consolidation. We then analyzed their long-term memory by exposure to the task 12 days later (Day 17, pink), observing that the TgQBP1 mice increased their escape latency compared to Day 5 (p=0.047) while the WT mice maintained a similar latency (**Fig. 3A**). This difference suggests that TgQBP1 mice remember worse platform location at the long-term, which indicates an impairment in memory consolidation. In fact, a new exposure on the next day (Day 18 –lilac-, **Fig. 3B**) showed a reduction in the escape latency of TgQBP1 mice in relation to the previous day (p=0.043), suggesting that TgQBP1 mice were then able to remember the platform. Together, these results suggest that TgQBP1 mice retain some memory (localization of the quadrant on Probe Q and decreased latency on Day 18), but have impaired long-term memory (failure to localize the platform on Probe P and increased latency on Day 17).

**Figure 3.**
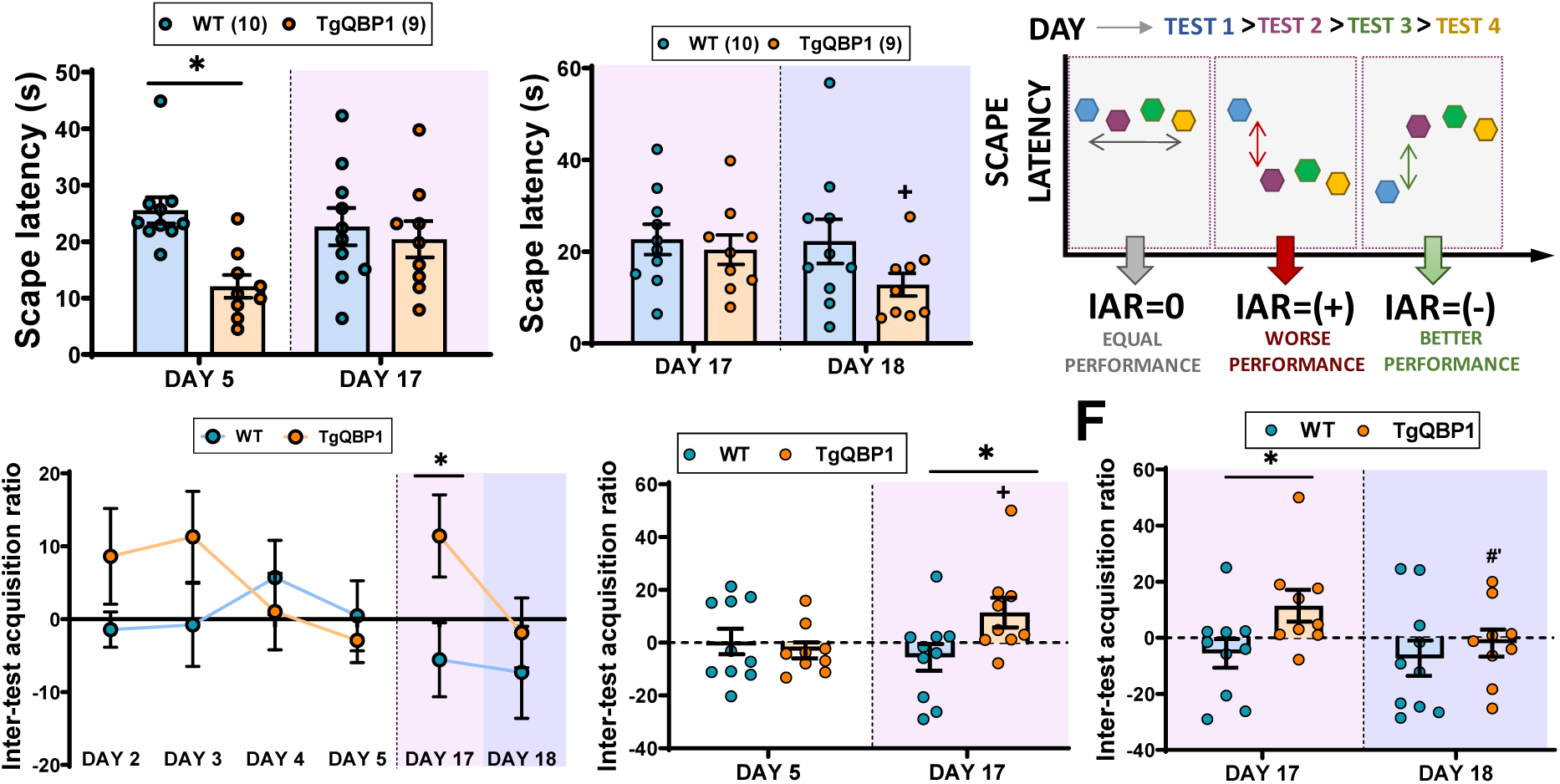
TgQBP1 mice showed intra-subject memory impairment in acquisition trials. **A)** Comparison of escape latency between Day 5 of acquisition and Day 17 (LTM, pink), showing a loss of significance of the last day of acquisition in comparison to the long-term test in TgQBP1 mice (p=0.047). **B)** Comparison of escape latency between Day 17 and Day 18 (LTM, lilac), showing an improvement of escape latency in TgQBP1 mice 24 h after LTM testing (p=0.043). **C)** Inter-test acquisition ratio (IAR) of the Morris water maze, which reflects the animal’s performance on the first test (Test 1) of each day of task acquisition. This ratio is obtained by subtracting the escape latency of Test 1 from the mean of the four tests performed each day of acquisition. If the animal’s performance on Test 1 is similar to the day’s mean (left panel), the IAR will have a value close to zero. On the other hand, if the animal’s performance is worse (middle panel), the IAR will be negative (higher escape latency in Test 1); while if the animal’s performance is better than the day’s mean (right panel), the IAR will be a negative value (lower escape latency in Test 1). **D)** Representation of the inter-test acquisition ratio (IAR) of WT mice (turquoise) relative to TgQBP1 mice (orange). The IAR of WT mice is close to zero in the first days of acquisition (equal performance in all tests) and then decreases as the mice remember the platform location (better performance). In the TgQBP1 mice, IAR is highest on the first days of acquisition (worse performance on Test 1 of each day) and then starts to decrease until day 5 (better performance because they remember the platform), but on day 17 it increases significantly (p=0.039; worse performance) and 24 h later it decreases again (better performance after re-exposure to the test). **E)** Comparison of the IAR between day 5 of acquisition and day 17 reflects that the TgQBP1 mice significantly worsen their performance on the first test on day 17 (p=0.042) with respect to the last day of acquisition. Also shown is the significant difference between groups on Day 17 (p=0.039). **F)** Comparison of IAR between Day 17 and Day 18 reflects that the TgQBP1 mice tend to significantly improve their performance on Test 1 on Day 18 with respect to Day 17 (p=0.091) and it has similar values to WT mice. For comparisons between independent groups: *p<0.05, **p<0.01, ***p<0.001, trends 0.05 ≥ # < 0.09; for comparisons between dependent groups: +p<0.05, ++p<0.01, +++p<0.001, trends 0.05 ≥ #’ < 0.09

To analyze this variation in performance, we analyzed whether impaired consolidation of TgQBP1 mice also occurs within each day of acquisition (24 h). Plotting the escape latency of the first test (Test 1) against the last test of the day (Test 4) showed that TgQBP1 mice significantly varied its performance along each test within the same day (**Fig. S3F**), specifically a trend on Day 5 (p=0.063) and a significant variation on Day 17 (p=0.040). In contrast, the performance of WT mice was homogeneous across all tests within a single day (**Fig. S3G**). We have defined a new variable we called Inter-test <u>A</u>cquisition <u>R</u>atio (IAR) to reflect the performance of the mice in Test 1 in comparison to the mean of the four tests of each acquisition day (**Fig. 3C**). IAR values for both groups reflected that WT mice had an IAR close to zero on the first days of acquisition (**Fig. 3D**, blue), which correlates with their performance being consistent across all tests on the same day. In contrast, TgQBP1 mice had very high IAR values along the first days of acquisition, suggesting a worse performance in the Test 1 compared to the other tests within a day (**Fig. 3D**, orange). In both groups, IAR value decreased from Day 4 onwards as they remember the task and improve their performance on Test 1. Finally, comparison of IAR values between acquisition (Day 5) and long-term re-exposure (Day 17, pink) reflected a significant increase of IAR value in the TgQBP1 mice relative to WT (p=0.039; **Fig. 3E**) and relative to their own performance on acquisition (p=0.042), again suggesting a worse performance after non-exposure to the task. However, this underperformance is recovered on Day 18 (lilac, **Fig. 3F**), where the IAR value of TgQBP1 mice was almost zero due to a tendency to improve their performance (p=0.091). Overall, the IAR variable has highlighted the impairment of long-term memory between each test day (every 24 h), evidenced by a worse performance on Test 1 that improves with subsequent test exposures within the same day. We suggest that, although QBP1 is inhibiting memory consolidation, the learning of this task is repetitive (4 test/day) and extended in time (3-5 days) so that memory consolidation seems to be good enough to recall the task after a single re-exposure (better performance on Day 18).

### QBP1 impairs long-term consolidation of aversive memories

After demonstrating that QBP1 was capable of impairing long-term consolidation of hippocampal-dependent memories, we proceeded to analyze aversive memories. To do this, we used the Contextual fear conditioning test, a paradigm of associative memory where the aim is to condition a traumatic event (small inescapable shock, unconditioned stimulus –UCS-) to a context and an acoustic cue (conditioned stimulus, CS) ^79^, following the experimental design described in **Figure 4A**. WT and TgQBP1 mice correctly acquired the aversive conditioning, which is reflected in the significant increase in the freezing percentage along each trial within a session (Trial p<0.001, **Fig. 4B**). After 24 hours, recall of the aversive context was analyzed by showing the freezing percentage per minute (**Fig. 4C**), where there was an interaction effect (p=0.008), based on both groups varying their behavior over time (Min p<0.001) and that TgQBP1 mice showed significantly less freezing than WT mice (Group p=0.046), suggesting impaired aversive memory in TgQBP1 mice.

**Figure 4:**
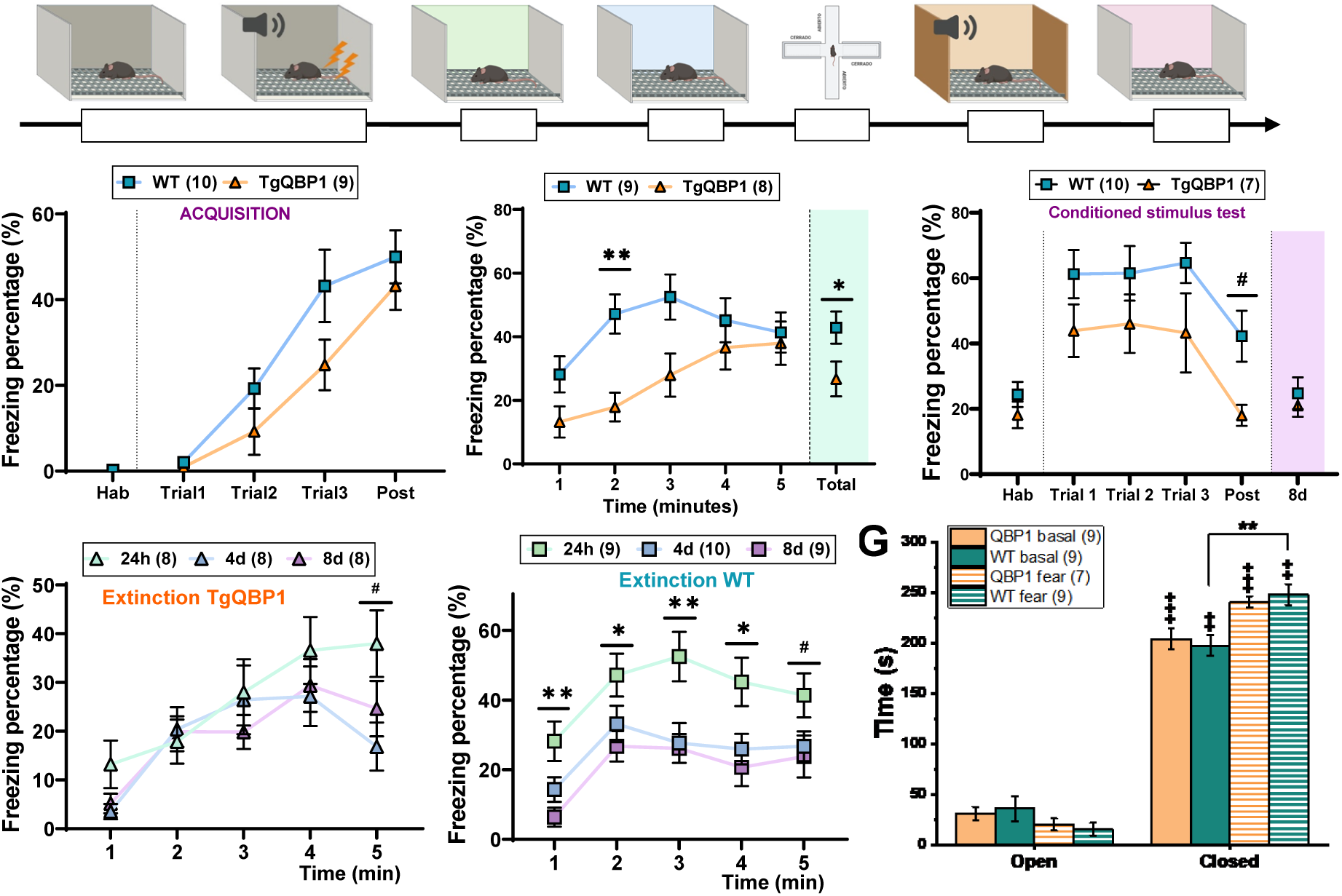
Analysis of aversive memories in TgQBP1 mice. **A)** Schematics of the experimental design for the contextual fear conditioning test. The diagram shows the temporal sequence of the test implemented in the TgQBP1 mice for fear conditioning implementation. **B)** Percentage of fear in the acquisition phase of fear conditioning, showing an increase in percentage with trials of the test for the two groups, with no significant differences between them. **C)** The percentage of fear in the context reviewed at 24 h after acquisition reflects that there are significant differences according to group and minute of the test (Interaction p=0.008). The TgQBP1 mice show a significant overall lower context fear behavior than WT (Group p=0.046), which broken down by minute shows a significant difference between groups at Min 2 (p=0.009). In addition, both groups show an overall increase in fear over test minutes (Time p<0.001). In terms of total time (green), the TgQBP1 mice showed significantly less freezing than the WT (p=0.046). **D)** The conditioned stimulus test (day 7) analyzes how the animal remembers the acoustic tone conditioned to the aversive event (shocks) in a different context (non-aversive). Both groups modify their behavior along trials (p=<0.001) and WT mice show a tendency to perform a higher percentage of global fear (Group p=0.051). This means that WT mice remember the conditioned stimulus, with a significant trend in the post-test phase with respect to the TgQBP1 mice (p=0.057). However, there are no differences in the total freezing percentage in context review 24 h after (8d). **E)** Intra-subject analysis of fear extinction per minute for the TgQBP1 mice. There are no differences in its overall behavior between the three days of context evaluation but it changes over the minutes of the test (24 h p<0.001, 4 d p<0.001 and 8 d p=0.03). Among extinction times, the TgQBP1 mice only tend to change its behavior at Min 5 (p=0.061). **F)** Intra-subject analysis of fear extinction per minute for WT mice. This group shows that animals change their behavior over the test minutes (p<0.001) and globally extinguish conditioning to context (p=0.002), especially from 24 h to 4 d-8 d (p<0.001). Furthermore, because of this, WT mice show different behavior per minute between days of exposure to context: Min 1 (p=0.005), Min 2 (0.044), Min 3 (p=0.008), Min 4 (p=0.016) and Min 5 (p=0.090). **G)** Elevated maze results to compare anxiety levels in basal *versus* aversive conditions. Total time spent in the arms showed that both groups and in all conditions spent significantly more time in the closed arms (TgQBP1: p<0.001 for basal condition and p<0.001 for aversive condition; WT: p=0.008 for basal and p=0.008 for aversive condition), but only WT mice significantly increased their time in the closed arms after fear conditioning (p=0.005). For comparisons between independent groups: *p<0.05, **p<0.01, ***p<0.001, trends 0.05 ≥ # < 0.09; for comparisons between dependent groups: +p<0.05, ++p<0.01, +++p<0.001, trends 0.05 ≥ #’ < 0.09

In the long-term, the recall of the conditioned stimulus was assessed (acoustic cue, **Fig. 4D**) and showed that both groups varied their behavior over trials (p<0.001) and TgQBP1 mice tended to exhibit overall less fear than WT mice (Group p=0.051). These results suggest that the TgQBP1 mice also showed impaired long-term memory of the conditioned stimulus. Finally, we compared the total freezing time on each aversive context test (24 h-4d-8d, **Fig. S4A**), reflecting that, although both groups reduced their freezing over days (Extinction p<0.001), only WT mice had a significant tendency to extinguish its freezing (p=0.006 and p=0.009 respectively). Overall, TgQBP1 mice showed a lower response to conditioned fear in both context and acoustic tone. According to our working hypothesis, this behavior may be attributed to an impairment in the consolidation process of the conditioned stimulus, leading to insufficient extinction of the memory. The analysis of aversive context recall within each group revealed differences in their behavior: TgQBP1 mice did not exhibit changes in overall behavior along days (**Fig. 4E**), indicating an absence of extinction due to impaired fear memory consolidation. However, within each day, their fear behavior significantly increased over the test minutes (24 h p<0.001, 4d p<0.001, and 8d p=0.03), suggesting that they recall the aversive context the more time they spend in it. In contrast, WT mice (**Fig. 4F**) displayed a linear and constant overall behavior and a global extinction of the aversive context over days (Extinction p=0.002). Indeed, there are differences among the three extinction tests along each minute of the test (**Fig. 4F**).

In order to correlate aversive memories with PTSD models, anxiety levels were evaluated on an Elevated plus maze after the fear conditioning, which were compared to levels in basal conditions (**Fig. S3 B-C**). In this comparison, both groups spent significantly more time in the closed arms, but only WT mice spent significantly more time in the closed arms after fear conditioning (p=0.005, **Fig. 4G**). This difference is related to the significant decrease in exploration of the open (p=0.001) and closed (p=0.005) arms of WT mice after the aversive conditioning (**Fig. S4B**), suggesting that this group hardly moves from the closed arms (protected). On the other hand, TgQBP1 mice explored significantly less in the closed arms (p=0.016). Furthermore, these differences in exploration do not correspond to animals predominantly entering some of the arms, as the percentage of entries did not reflect any significant difference (**Fig. S4C**). In conclusion, TgQBP1 mice do not show a significant increase in anxiety levels after the traumatic event, unlike WT mice. This indicates that QBP1 not only impairs memory consolidation but also reduces the anxiety induced by aversive events.

### Effects of age and sex on memory impairment in TgQBP1 mice

After demonstrating the effect of QBP1 on memory consolidation in adult male mice, we decided to expand our study to investigate the effects of age and sex. Firstly, it has been reported that younger individuals develop PTSD more often due to their more vulnerable brains and greater exposure to high-risk situations ^10,80^. This can significantly impact their lives for over a decade and impose a substantial economic burden on the healthcare system ^80,81^. Also, aversive memory consolidation in adolescent mice differ in extinction compared to adults, which may explain the challenges in treating anxiety disorders in younger populations ^81,82^. To analyze the effect of QBP1 in young animals, male mice aged 2-3 months were subjected to contextual fear conditioning (**Fig. 5A**). During acquisition, TgQBP1 and WT groups significantly learned the task (Trial p<0,001) with no differences between them. Analysis of the aversive context recall at 24-96 h indicated that both groups globally extinguished freezing behavior (Time p=0,003) but showed overall differences (Group p=0,054), with young TgQBP1 mice showing a trend to perform less total freezing compared to WT mice at 24 h (p=0,051). This suggests that the effect of QBP1 on impaired aversive memory consolidation is similar in young male mice.

**Figure 5.**
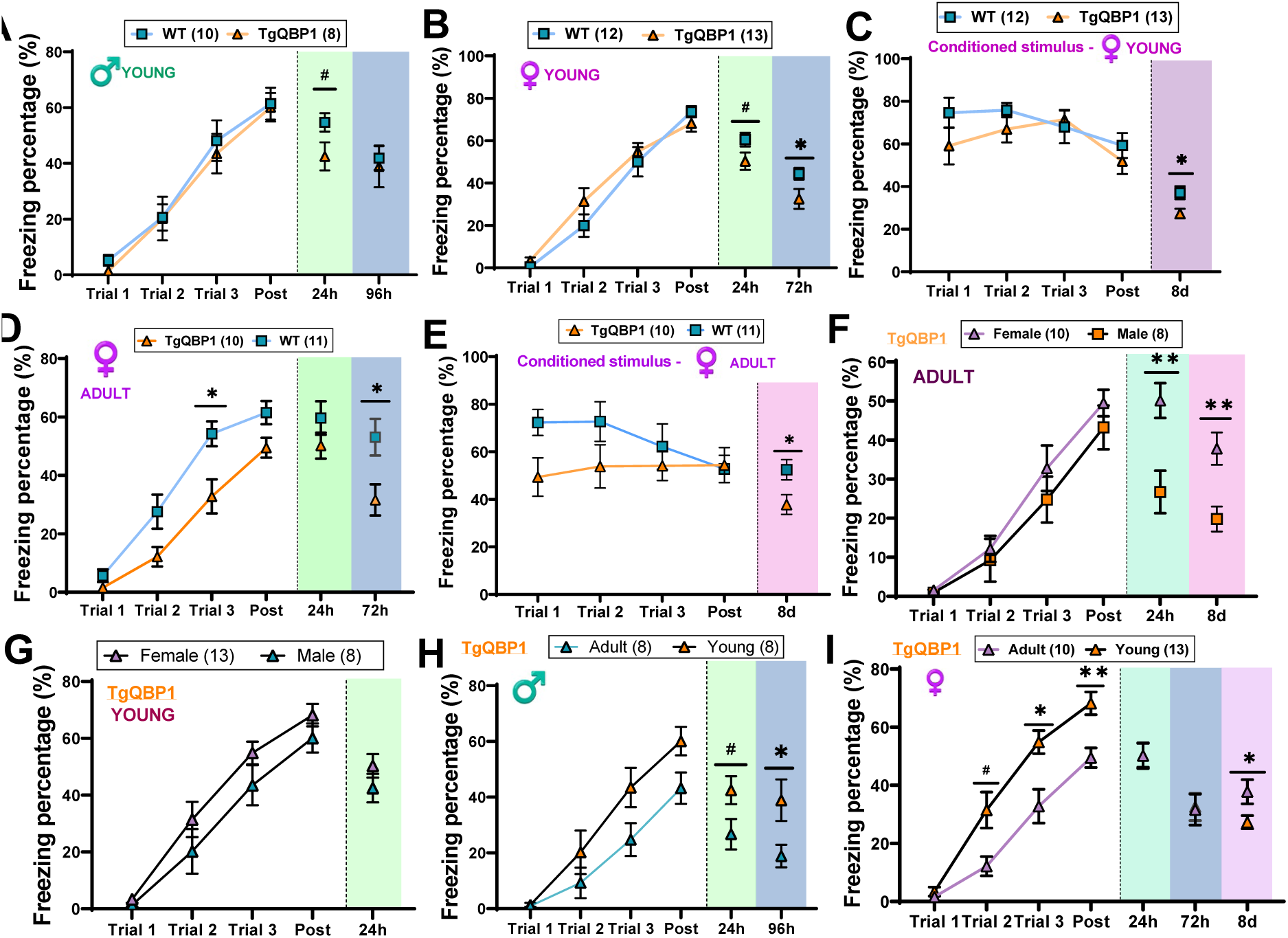
Effects of Age and Sex on Fear Conditioning in TgQBP1 Mice. **A)** Percentage of fear in young male mice during aversive conditioning. During the acquisition phase, both groups learned the conditioning task significantly with trials, with no differences between them. However, at 24 h post-training, TgQBP1 mice showed a trend towards poorer recall of the fear context compared to WT mice (p=0.051). By 96 h, this significance of this difference was lost as both groups exhibited similar fear extinction. **B)** Percentage of fear in young female mice during aversive conditioning. Both groups learned the conditioning task significantly during acquisition, with no differences between them. However, at 24 h post-training, TgQBP1 mice showed a trend towards less total fear compared to WT mice (p=0.066), with this difference becoming more significant at 72 h (p=0.044). **C)** Percentage of fear in young female mice during conditioned stimulus recall (acoustic, different context). there were no significant differences between the two groups during the acoustic stimulus trials. However, at 24 h post-training (8 days after acquisition), WT mice exhibited significantly more total fear than TgQBP1 mice (p=0.016). **D)** Percentage of fear in adult female mice during aversive conditioning. Both groups learned the conditioning task significantly during acquisition (p<0.001), with TgQBP1 females showing significantly less global fear compared to WT mice (p<0.001). However, at 24 h post-training, no differences were observed between the groups, while at 72 h, TgQBP1 mice performed significantly worse than WT mice (p=0.018). **E)** Percentage of fear in adult female mice during conditioned stimulus recall (acoustic, different context). No significant differences were observed between the groups during the acoustic stimulus trials. However, at 24 h post-training (8 days after acquisition), TgQBP1 mice performed significantly worse than WT mice (p=0.023). **F)** Effect of sex in adult TgQBP1 mice during aversive conditioning. During acquisition, both groups learned the conditioning task significantly with trials, with no differences between them. However, at 24 h post-training, adult males showed significantly poorer recall of the fear context compared to adult females (p=0.003), which persisted at 8 days (p=0.004). **G)** Effect of sex in young TgQBP1 mice during aversive conditioning. During acquisition, both groups learned the conditioning task significantly with trials, with no differences between them. This lack of differences persisted at 24 h post-training, indicating no gender differences in young animals. **H)** Effect of age on aversive memory consolidation in male TgQBP1 mice. Both age groups learned the conditioning task significantly during acquisition (p<0.001), with a significant overall age effect (p=0.044). This effect persisted at 24 h and 96 h post-training, with young TgQBP1 mice exhibiting significantly more fear than adults (p=0.052 and p=0.033, respectively). **I)** Effect of age on aversive memory consolidation in female TgQBP1 mice. Both age groups learned the conditioning task significantly during acquisition (p<0.001). However, young females exhibited significantly more fear than adults during acquisition (interaction p=0.024 and age p<0.001), particularly in Trial 2 (p=0.054), Trial 3 (p=0.025), and Post (p=0.007). Post-training context review showed that adult females exhibited significantly less fear than young females at 8 days (p=0.028), indicating greater long-term memory consolidation impairment in young females. For comparisons between independent groups: *p<0.05, **p<0.01, ***p<0.001, trends 0.05 ≥ # < 0.09; for comparisons between dependent groups: +p<0.05, ++p<0.01, +++p<0.001, trends 0.05 ≥ #’ < 0.09

Secondly, considering that women are twice as likely to develop PTSD ^10,83,84^, we decided to analyze the consolidation of aversive memory in female mice. These differences may be related to the predominance of the basal amygdala in fear consolidation in females, who tend to exhibit greater fear response to context and generalized fear ^85–87^. For our study, female mice aged 2-3 months were exposed to contextual fear conditioning, with context revision performed at 24-72 hours considering that females exhibit a faster loss of specific context memory ^87^. Acquisition indicated that both groups properly learn the aversive conditioning (Trials p<0,001) without differences between them (**Fig. 5B**). The total freezing percentage in aversive context recall (24-72 h) showed that young TgQBP1 females tended to exhibit significantly less freezing at 24 h compared to WT mice (p=0,066), a difference that remained significant over time (72 h p=0,044). Furthermore, analysis of the conditioned stimulus (acoustic cue, **Fig. 5C**) revealed no significant differences between the groups, but the analysis of aversive context recall after the stimulus in young female mice (8 days post-acquisition) maintained the significant difference between groups (p=0,016). These results confirmed the effect of QBP1 on memory consolidation in young female mice. Similarly, we analyzed the stability of aversive memories in adult female mice (**Fig. 5D**). Both groups significantly learned the conditioning during acquisition (Trials p<0.001), but adult TgQBP1 females exhibited a significantly lower overall freezing behavior during acquisition compared to WT mice (Group p<0.001), with a significant difference in Trial 3 (p=0.034). However, the analysis of aversive context recall at 24 hours showed no differences between groups, while the adult TgQBP1 females displayed significantly less total fear than WT mice at 72 hours (p=0.018; **Fig. 5D**, blue). Lastly, analysis of the conditioned stimulus recall in adult female mice (auditory cue) showed no significant differences between groups (**Fig. 5E**), despite WT mice performed higher freezing percentages. A significant difference was observed in the context review at 8 days, where adult TgQBP1 females showed significantly lower total freezing percentages compared to WT mice (p=0.023). Together, these data indicate that the impairment in aversive memory consolidation is also present in adult female TgQBP1 mice.

Next, a comparative analysis of all behavioral data was conducted to draw conclusions regarding the effect of age and sex on the deterioration of aversive memory in TgQBP1 mice. First, the effect of gender was analyzed by comparing each age group separately (**Fig. 5F-G**). In adults, both sexes learned the aversive conditioning significantly without differences between them (Trial p<0.001; **Fig. 5F**). However, adult females showed significantly better recall of the aversive context compared to adult males at 24 hours (p=0.003) and 8 days (p=0.004). In young mice, both sexes also learned the aversive conditioning without significant differences between them (Trial p<0.001; **Fig. 5G**), and this lack of difference persisted in the 24-hour context review. Together, these results suggest that although the ability to learn aversive conditioning is similar across genders and ages, adult females exhibit better retention of aversive memory compared to adult males, a difference not observed in young mice.

Furthermore, the effect of age was analyzed, separately comparing each gender (**Fig. 5H-I**). In male TgQBP1 mice, the acquisition reflected that both age groups significantly learned the conditioning (Trials p<0.001), with an age effect observed (Age p=0.044; **Fig. 5H**). This significant difference persisted in the analysis of aversive context recall at 24 h (p=0.052) and 96 h (p=0.033), where adult mice showed significantly less fear than younger mice. Regarding female TgQBP1 mice (**Fig. 5I**), the conditioning acquisition showed a significant interaction between variables (p=0.024), indicating both age groups significantly learned the conditioning (Trials p<0.001) and there was an age effect (Age p<0.001). Specifically, younger females exhibited significantly higher freezing percentages than adult females during acquisition phases Trial 2 (p=0.054), Trial 3 (p=0.025), and Post (p=0.007). However, during aversive contextual recall analysis, no differences between age groups were observed at 24-72 h, but younger females exhibited significantly less fear than adults at 8d (p=0.028). In conclusion, age is suggested to have a significant impact on aversive memory recall in the TgQBP1 mice of both sexes. Specifically, younger mice tend to show greater freezing behavior during acquisition. However, this effect was reversed during long-term context reviews, where young males showed higher freezing percentages.

Along these lines, the analysis of WT mice showed a behavior similar to that of TgQBP1 mice in terms of age and sex (**Fig. S5**). Adult female mice showed significantly more freezing in the analysis of aversive context recall at 24 h and 8d compared to adult males (p=0.045 and p=0.009 respectively, **Fig. S5A**), whereas young mice did not show sex differences (**Fig. S5B**). Regarding age, young WT males showed significantly more freezing than adults in the aversive context analysis at 24-96 h (p=0.061 and p=0.021 respectively, **Fig. S5C**), while no significant age-related differences were observed in WT females except for a significant freezing increased at 8d (p=0,004, **Fig. S5D**). Taken together, the behavior of WT mice is similar to that of TgQBP1 mice, demonstrating that age and sex matter when analyzing memory consolidation, but the presence of QBP1 in the transgenic mice did not promote additional differences.

### mCPEB3 showed reduced aggregation in TgQBP1 mice

Once the effect of QBP1 on mouse memory consolidation was demonstrated, we wanted to study its mechanism of action based on our working hypothesis, *i.e.* that QBP1 inhibits the amyloidogenesis of CPEB3 and that this blockade is what causes memory impairment. After performing the memory tests, the hippocampi of the mice tested were extracted and mCPEB3 was characterized following a detergent insolubility protocol specific for amyloid proteins ^58,60^, assuming that, after undergoing a learning process, mCPEB3 oligomerizes and increases its presence in the insoluble fractions ^56,60^. This protocol uses strong detergents in the lysis buffer and ultracentrifugation to separate the cytosolic material into two fractions (**Fig. S6A**): aggregated protein (amyloid, precipitate P2) or soluble protein (supernatant S2). Western blot analysis of the band corresponding to mCPEB3 (75 kDa) was then performed comparing basal conditions of WT mice (WT naive, **Fig. 6A**) with the conditions after learning in WT and TgQBP1 mice (WT learn and TgQBP1 learn, **Fig. 6A**), comparing the soluble (S2, soluble protein) and precipitated (P2, aggregated protein) fractions corresponding to the cytosol fraction.

**Figure 6:**
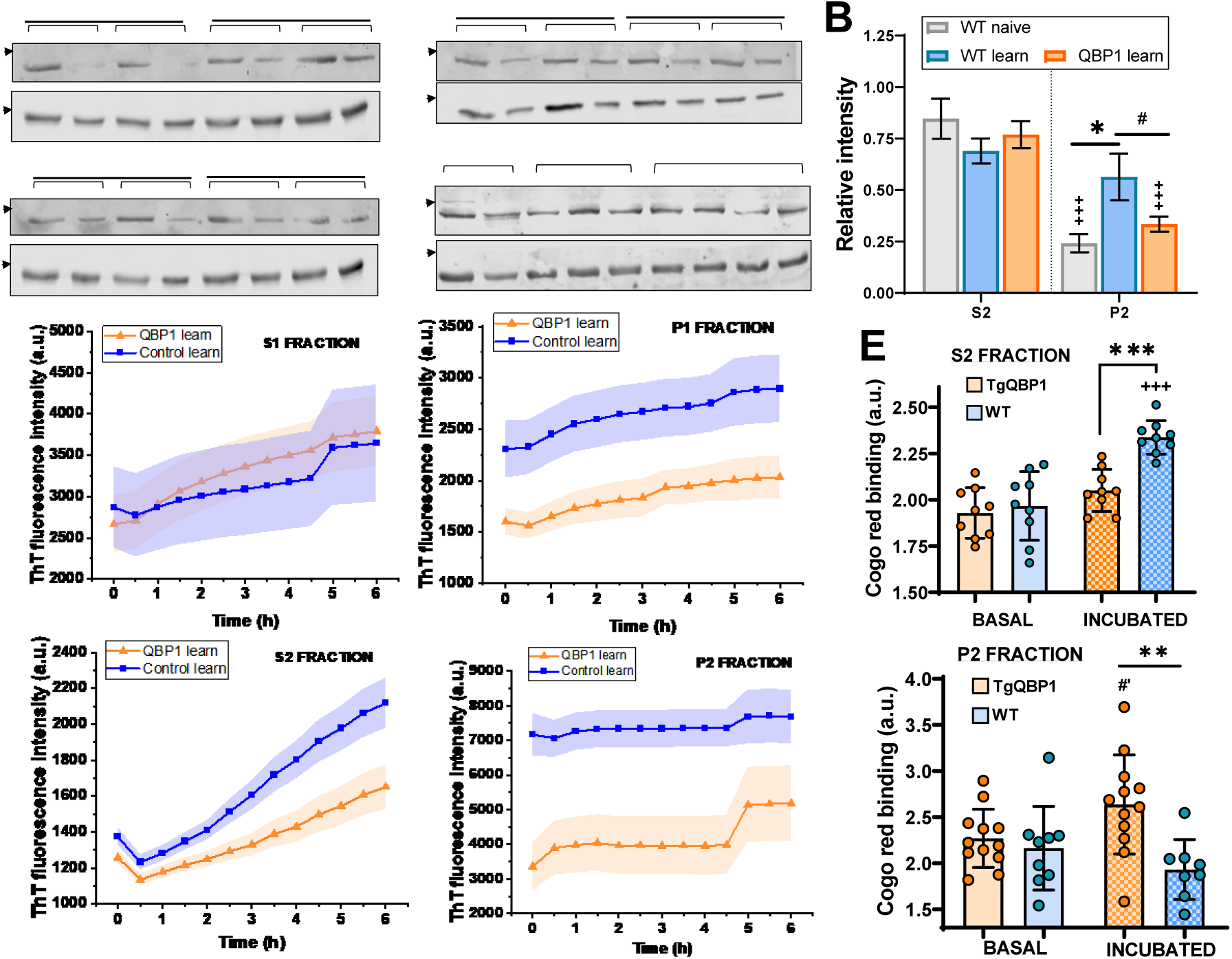
Analysis of mCPEB3 protein in TgQBP1 mice (learning *versus* basal conditions). **A)** Representative images of Western blots from the analysis of soluble (S2) and precipitated (P2) fractions of hippocampal samples, comparing the WT naive group versus a learning conditions of both groups for mCPEB3. GAPDH was used as a loading control. **B)** Relative intensity corresponding to the quantification of the 75kDa band in the S2 and P2 fractions of mCPEB3 in hippocampi (Image J software, N=5 animals/group). The values of WT mice under naive conditions (WT naive, gray) were compared to those after an aversive learning corresponding to both WT (blue) and TgQBP1 (orange) groups. Statistical analysis revealed that there is an interaction effect between variables (Fraction*Group=0.008) and in the Fraction variable (p<0.001), which is reflected in the fact that P2 fraction is significantly elevated in the WT learn relative to the WT naive (p=0.016) and in a trend in comparison to TgQBP1 learn (p= 0.080). At the intrasubject level, WT naive and TgQBP1 fear groups significantly change their relative intensity between S2-P2 fractions (p>0.001), which is not the case for WT fear. **C)** Aggregation kinetics of the fractions extracted from hippocampal S1-P1 after learning monitored by Thioflavin T fluorescence intensity. The values corresponding to the first fractions of the extraction plotted in the graphs, showing that, while there is hardly any difference in aggregation in the cytosolic fraction (S1) there are clear differences in the insoluble fraction (P1). **D)** Aggregation kinetics of the cytosolic fractions extracted from the hippocampi (S2-P2) after learning monitored by Thioflavin T fluorescence intensity. In both cases, the formation of mature amyloid over time is higher for WT mice, particularly in the insoluble fraction (P2) where there is a higher proportion of aggregated protein. **E)** Amyloid quantification based on Congo Red (CR) binding in the S2 and P2 fraction of hippocampi extracted after learning. CR binding of the soluble fraction (S2, upper panel) reflects that there is interaction (p=0.024), where the effect of incubation and group are significant (p<0.001 both). In fact, the WT sample significantly increases its value after incubation (p<0.001) and with respect to the TgQBP1 sample (p<0.001). The binding to CR of the insoluble fraction (P2, lower panel) shows that only the TgQBP1 sample tends to increase its value with incubation (p=0.067) and differs significantly from the WT sample (p=0.002). For comparisons between independent groups: *p<0.05, **p<0.01, ***p<0.001, trends 0.05 ≥ # < 0.09; for comparisons between dependent groups: +p<0.05, ++p<0.01, +++p<0.001, trends 0.05 ≥ #’ < 0.09

To analyze possible differences between the groups, quantification of the band corresponding to ∼75 kDa (monomeric canonical form of mCPEB3) was performed in all animal samples analyzed for fractions S2 and P2 (**Fig. 6B**), obtaining the relative intensity value referred to its loading control (GAPDH) for each extraction (Image J software). The analysis reflects that the relative intensity with respect to the mCPEB3 band in the soluble fraction (S2) is similar between groups, with no differences between the WT naive (gray) and both groups after aversive learning (WT learn –blue- and TgQBP1 learn –orange-, **Fig. 6B left**). However, the relative intensity in the aggregated fraction (P2) in the WT learn is significantly increased with respect to the WT naive values (p=0.016) and shows a trend in the TgBQP1 learn group (p=0.080) (**Fig. 6B right**). These results show that the insoluble fraction increases when the animal experiences learning, correlated to the aggregation of the mCPEB3 protein needed to perform its function ^60^. On the other hand, the presence of QBP1 in the transgenic mice causes an impairment of long-term memory that is accompanied by a reduction of protein aggregation, similar to the values of WT mice without learning (WT naive). Taken together, these results suggest that the molecular mechanism underlying the long-term memory impairment found in the TgQBP1 mice could indeed correspond to a decreased mCPEB3 oligomerization produced by the presence of QBP1.

### QBP1 reduces mCPEB3 amyloid state in TgQBP1 mice after learning

In order to analyze the amyloid properties of CPEB3 in mice brain, hippocampal samples isolated and extracted after learning were characterized. Firstly, amyloid propensity was characterized using ThT fluorescence aggregation kinetics, where higher values indicate greater aggregation potential. The results indicate minimal differences in the cytosolic fraction (S1), but in the insoluble fraction (P1), the WT sample aggregated more than that from the TgQBP1 mice (**Fig. 6C**). Furthermore, this difference was maintained in the S2 (soluble protein) and P2 (aggregated protein) fractions from animals that underwent learning (**Fig. 6D**), suggesting that samples from WT mice show greater amyloidogenic potential and aggregation compared to samples where QBP1 is present.

Next, the aggregation potential was evaluated through its binding to Congo Red, a specific staining characterized by a shift in its absorption peak from 490 to 512 nm and an additional peak at 540 nm upon its binding to the mature amyloid structure ^88,89^. In this case, an additional incubation (6 h at 37 °C) was performed to evince the amyloid potential of the sample. After binding quantification (**Fig. 6E**), S2 fraction (soluble protein) showed that the sample from WT mice significantly increased its binding after incubation (p<0.001) compared to the TgQBP1 mice sample (p<0.001). The P2 fraction (aggregated protein, lower panel) exhibited fewer differences between groups, with the TgQBP1 sample tended to increase after incubation (p=0.067) and significantly changed in correlation with the WT incubated sample (p=0.002). Taken together, these results suggest that the samples with the highest amyloid potential are the S2 fraction from WT mice and the P2 fraction from TgQBP1 mice, indicating that they contain the highest proportion of protein able to aggregate. The fact that P2 fraction (aggregated protein) of TgBQP1 mice maintain an increased amyloid potential suggests that the presence of QBP1 has blocked its amyloid conversion.

Finally, the insoluble fractions (P1-P2) of the learned samples were characterized using immunodot blot against A11/OC conformational antibodies, with CPEB3 as a control (**Fig. S6B**), followed by quantification using Image J (**Fig. S6C**). Quantification did not reveal any significant differences between groups in A11/OC reactivity, although differences were observed across fractions (A11 p=0.005 and OC p=0.002). However, CPEB3 reactivity indicated no interaction effect but showed significant group and fraction effects (p<0.001 both), with TgQBP1 mice displaying a significant increase in P1 reactivity (p=0.007). This difference suggests that mCPEB3 in TgQBP1 samples is less aggregated and more accessible to antibody interaction within insoluble fractions. Taken together, this biochemical and amyloid characterization suggests that the presence of QBP1 in the TgQBP1 mice is decreasing the amyloid potential of CPEB3, evincing that the mechanism of action of QBP1 involves the amyloid blockade of the protein.

## Discussion

Considering the relevance of the amyloid state of CPEB3 in memory consolidation ^47,60,90^ and the tight control that exists over its functional conversion ^58,59,91^, this protein stands as a molecular switch of long-term memory consolidation. We already proposed CPEB *Drosophila* orthologue, Orb2, as a therapeutic target to impair memory consolidation ^74^ and as a potential approach towards PTSD treatment, as this disorder is caused by the poor adaptation of the brain to a traumatic memory ^26,92^. Considering that PTSD current pharmacology is mostly based on reducing the associated symptomatology ^17,93,94^, the QBP1 peptide^65,95^ was chosen as a potential compound for a pharmacological intervention as it has already been shown to be effective against other amyloid proteins with polyQ regions by blocking their amyloid conversion of their monomer ^69,70,96^.

We have shown that the QBP1-M8 peptide effectively inhibits the amyloid state of the human CPEB3 protein. Specifically, it inhibits the aggregation kinetics of the IDR region and its amyloid core (residues 101-200) at ratios 1:4 and 1:8, similar to TDP-43 ^70^ and Orb2A ^74^. Notably, TEM visualization reveals that the interaction of QBP1-M8 with both CPEB3 and Orb2A^74^ leads to an increase in nonspecific aggregates rather than the formation of mature amyloid fibers. However, QBP1-M8 effectively inhibits the amyloid core fibrillation (residues 101-200), but it does not inhibit the entire amyloid-forming subdomain (residues 1-200) (**Fig. 2**). According to this, the QBP1-M8 effect could depend on the structure adopted by each fragment. On one hand, the amyloid core is composed by a hydrophobic region with rigid β-sheet-rich segments ^97–99^ and a Pro/Ala/Gln-rich region similar to the region characterized in TDP-43 (341-357) ^70^. This means that their glutamines would remain accessible for interaction with QBP1 ^70,100^. On the other hand, the amyloid subdomain (R2_1-200_) is composed by two long Pro/Ala/Gln-rich regions flanking the hydrophobic region, whose organization into different α-helices would end up promoting its amyloid aggregation ^99^, leading to the acquisition of a folded structure inaccessible to QBP1-M8. Related to this, we have showed that SCR-C peptide (QBP1-M8 random sequence) is also capable to inhibit the amyloid kinetics of hCPEB3 IDR, as occurs in TDP-43 ^70^ and contrary to what occurs with Orb2A and *Ap*CPEB ^74,101^. Considering that the IDRs of TDP-43 and CPEB3 are based on short Q/N-rich regions, whereas those of *Ap*CPEB and Orb2 are mostly composed by long glutamine tracts ^70,72,75,101^, we suggest that the polyvalence and affinity of QBP1-M8 inhibition may lie on its structure (hydrophobic pocket and tryptophan arrangement) ^72^, while SCR-C mechanism may depend on the specific interaction between its tryptophan and asparagine residues and other hydrophobic residues present on the Q/N-rich IDRs. The current literature supports this hypothesis, where the development of novel anti-amyloidogenic peptides against the low complexity domain of TDP-43^102^ suggests that the aromatic cluster W(x)WW(x) (residues shared by QBP1-M8, SCR-C and 1b2-30-alt6 peptides) could be key in increasing the specific interaction with proteins rich in Q/N regions.

The generation and characterization of a transgenic mouse constitutively expressing QBP1 reported in this study constitutes the first proof-of-concept in mammals that this peptide is a good candidate as a therapeutic approach for PTSD. Phenotypic characterization of TgQBP1 mice has revealed normal function and development, with a correct neurological profile. However, analysis of memory consolidation has evidenced several long-term impairments, consistent with previous results in *Drosophila* ^74,101,103^. Specifically, TgQBP1 mice significantly decreased memory consolidation at 24 h in short learning (activity cage, object recognition and fear conditioning) while it maintained memory in repeated learning (Morris water maze) with the impairment occurring in the first trial of the first days of acquisition (IAR, **Fig. 4**). These results highlight that the type and number of exposures of each learning influence the reinforcement of memory consolidation ^104^, that is, the repetition of a learning could correspond to an increase in the number of times that CPEB3 adopts its amyloid state ^60,105,106^, thus favoring an increase in its accumulation since its prion-like characteristics promote its stability over time ^56,98^. Thus, each day of acquisition in the Morris water maze increases the amount of amyloid CPEB3, thus limiting the action of QBP1 and promoting proper long-term memory consolidation. Likewise, it is noteworthy that TgQBP1 mice have limited fear extinction capacity (**Fig. 5**), which may suggest a rigidity of synaptic plasticity ^41,107,108^.

We have then examined the mechanism of action underlying the blockade of memory consolidation in the TgQBP1 mice by analyzing the mCPEB3 protein as a therapeutic target for the QBP1 peptide. In hippocampal samples, the band corresponding to mCPEB3 (75 kDa) ^51,109,110^ reflects an increase in the fractions corresponding to protein aggregated in the cytosol (P2) only in WT mice after an aversive learning (WT learn). On the contrary, values of TgQBP1 mice after learning remain similar to those of WT mice without learning (WT naive). Subsequent examination of the amyloid potential within samples derived from both groups evidenced that WT mice samples aggregated significantly more than TgQBP1 mice samples, showing the underlying inhibition of the QBP1 peptide on the amyloid state of the protein. Taken together, these results confirm that mCPEB3 increases its amyloid and aggregated state after learning ^60^, but the presence of QBP1 limits the acquisition of this active amyloid state in the TgQBP1 mice, which in turn impairs memory consolidation.

Moreover, our results showed that QBP1 is capable of reducing memory consolidation independently of sex and age, since TgQBP1 mice showed a long-term memory impairment in each analyzed condition. However, it is worth mentioning certain differences in behavioral responses regarding sex and age, although they are also present in WT mice too. Our results have shown that both WT and TgQBP1 adult females maintain freezing levels significantly higher than those of males in the context text, while no significant differences are observed in younger animals. Current studies have described similar sex differences for fear responses ^86,111^, mostly related to the increased amygdala activation that underlies fear responses in females as opposed to the shift toward hippocampal activation made by males during the expression of contextual fear ^83,112^. In terms of age, impaired memory consolidation appears significantly more pronounced in young animals compared to adults, potentially due to an age-related compensatory mechanism. This compensation could be explained by the cumulative activation of cellular machinery each time a memory consolidation attempt occurs. Such repeated activation may promote transient engagement of synaptic plasticity pathways ^113,114^, although these mechanisms typically do not persist over time in the absence of consolidation-associated stimuli (e.g., “labels” or synaptic tags) ^113,115^. Over time, processes like synaptic plasticity and the formation of new synaptic contacts tend to diminish, particularly as learning extinction progresses^116^. However, events independent of de novo protein synthesis, such as intracellular signaling or cytoskeletal remodeling, may persist ^117^. These lasting molecular changes are the ones that could facilitate a form of sensitization or compensatory plasticity in older animals, ultimately mitigating the impact of memory consolidation impairments in comparison to younger animals.

As an important derivative of these results, the TgQBP1 mice represents as an interesting opportunity to clarify the current controversy on the role of CPEB3: for some authors, CPEB3 is considered a repressor of protein synthesis and its blockade would enhance memory consolidation ^50,90,118^, while for others it is an activator of protein synthesis and its blockade would impaired memory consolidation ^58–60^. **Figure 7** lists all existing mice models described in the literature related to CPEB3 modulation. When CPEB3 expression is abolished (KO CPEB3) ^90^, the effect on the phenotype relies on the absence of its repressor state, thus resulting in improved learning and an enhanced long-term memory. Conversely, if CPEB3 expression is specifically abolished after learning (cKO CPEB3) ^60^, the phenotype relies in the absence of its active state, which results in an impairments in 24-h memory retention and a spatial cognitive deficit in long-term platform localization. Finally, when the positive post-translational modification (Neurl1DN mice model) ^59^ or the amyloidogenic state of CPEB3 (TgQBP1) is modulated, the active state of CPEB3 is blocked and promotes impaired long-term memory and reduced cognitive flexibility and spatial discrimination. Taken together, we suggest that the properties of its active state are the ones that determine its effect on memory consolidation ^91,105^, rather the presence of the protein itself. Therefore, we conclude that CPEB3 appears to be a molecular switch for memory consolidation that exerts a dual function depending on its conformational state, where both states are equally relevant for memory consolidation and synaptic plasticity associated with learning.

**Figure 7:**
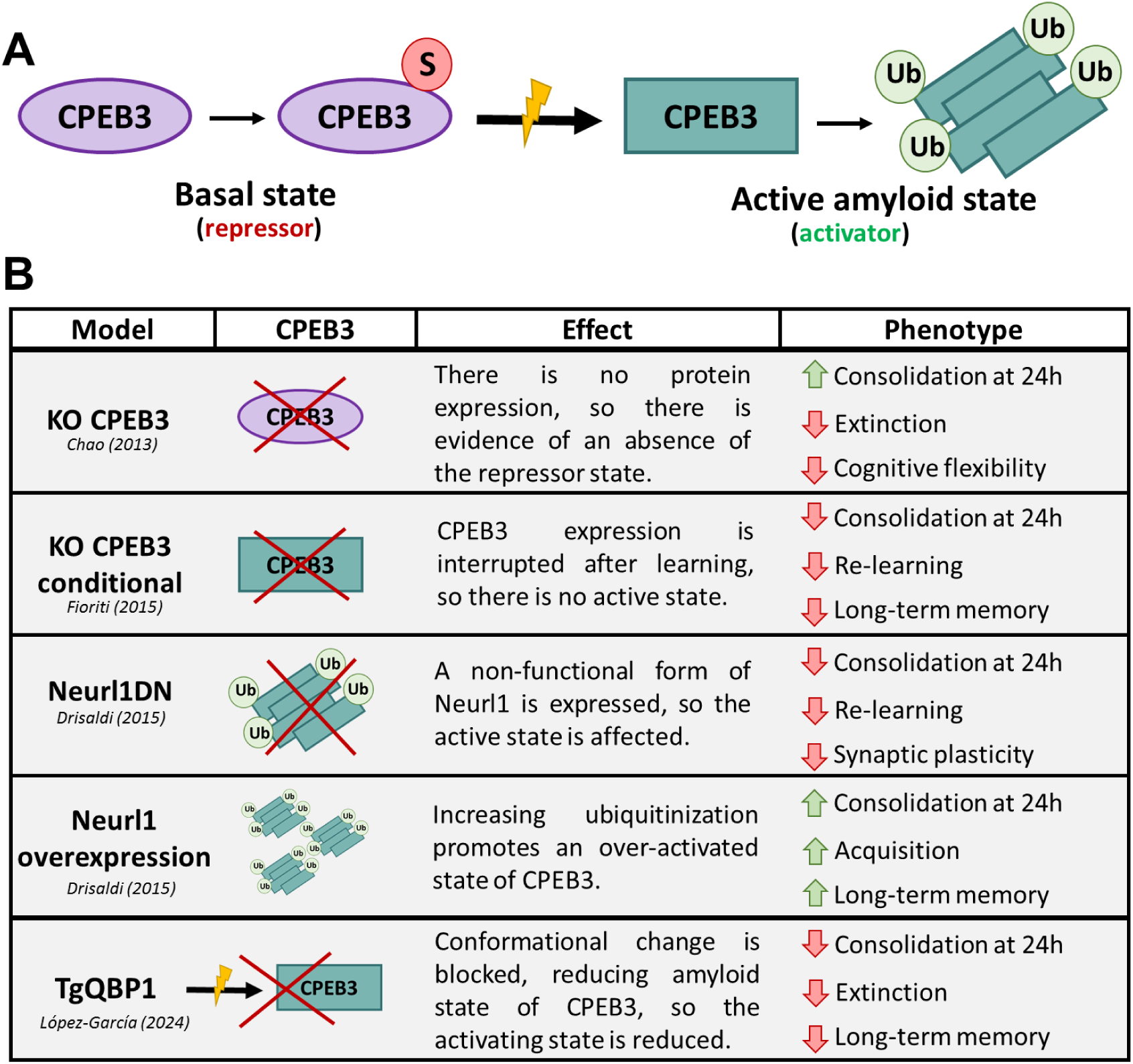
Comparison of the state of the CPEB3 protein modulation in mice models. **A)** Schematics of the CPEB3 conformational change, illustrating the amyloidogenic modulation of the protein depending on its state. CPEB3 is mostly found on its repressive monomeric state in basal conditions, with a sumoylation as a constraint that, after synaptic stimulation, give raise to the prion-like/amyloidogenic state that is ubiquitinated (positive posttranslational modification) and acts as an activator of synaptic plasticity by promoting mRNAs translation. **B)** Table summarizing the CPEB3 mice models found in the literature, indicating the modification involved on the CPEB3 protein and the repressor/activator function that is altered, which is directly reflected in a mice memory phenotype.

Regarding PTSD, the literature has recently established *cpeb3* as a potential risk gene for PTSD after evaluating the exacerbated and non-extinguishable fear response in the CPEB3 KO mice ^50^, which was associated to an increase in glucocorticoid rector levels and BDNF depletion in these animals. Since this is a model that completely lacks of CPEB3 protein ^90,118^ and only a phenotype where there is no basal CPEB3 repressor state is performed (**Fig. 7**), the increased susceptibility to fear may actually be due to the potentiation of memory acquisition that arises in this model and which occurs also for tasks where no aversive memories are involved. Our results on TgQBP1 mice show that the defect in memory consolidation is accompanied by anxiety levels similar to their baseline conditions (**Fig. 5G**), which suggests that QBP1 peptide not only decreases memory consolidation (by blocking CPEB3) but also reduces the negative emotional impact of the aversive event. Taken together, both studies demonstrate that CPEB3 is involved in memory consolidation and that, depending on how learning occurs and under what conditions ^50,60,90,104^, these memories will have a greater or lesser emotional impact ^26,32,119^.

Concerning the future preclinical development of QBP1 for PTSD, our results from the TgQBP1 mice suggest that blocking long-term memory provides an emotional benefit that reduces anxiety in such animals to baseline-like conditions, which suggests that QBP1 can be considered a causal treatment for the disorder which addresses this essential need in the current pharmacological treatment of PTSD ^23,46,120^. If we look at the current pharmacology based on fear memory ^26,42,119^, propranolol might be the most comparable drug to QBP1, since it affects the reconsolidation process by indirectly blocking protein synthesis ^42,121^. In mice, the effects of propranolol on both contextual and fear memories have been demonstrated ^121–123^, but unfortunately it has failed to prevent PTSD-like in rats ^121,124^ and has shown mixed results in human trials ^6,43,93,125^. On the other hand, hydrocortisone has shown beneficial effects in both rodents and humans ^37,38,41,93^, which makes it the most promising fear-based drug for the treatment of PTSD today. Considering that the behavioral protocols used in mice to evaluate the effects of propranolol and hydrocortisone ^121,124,126^ are similar to those we have used here for TgQBP1 mice, we can conclude that the positive results obtained with this peptide are comparable to those obtained by these current alternative drugs. Thus, QBP1 constitutes a promising alternative showing an effect comparable to that of propranolol and hydrocortisone, with the advantage that it affects a direct target that interferes with protein synthesis (CPEB3) to disrupt memory consolidation instead of just modulating the emotional response associated to such memory (*i.e.*, stress response axis disruption).

## Conclusions

At the level of basic research, our study has directly demonstrated the amyloidogenic hypothesis for neuronal CPEB in a mammalian system; while, at the level of biomedical research, we have provided the proof of concept that the anti-amyloidogenic QBP1 peptide can block memory consolidation in mice, showing therefore its effectiveness in mammals. Thus, QBP1 constitutes a new pharmacological tool that is very promising for the development of new therapies based on amyloid inhibition of CPEB. In addition to showing the safety of the peptide through its constitutive expression, we have demonstrated how QBP1 affects the new contextual and aversive memories, which points to QBP1 as a future strong alternative to hydrocortisone and propranolol in fear-based approaches for PTSD.

## Materials and methods

### Production of the QBP1 construct

The insert expressing two copies of QBP1 (in tandem) fused to eGFP was cloned into the pCAGGS expression vector, which contains the CAG promoter into the rabbit *β*-globin splicing intron and which ended with polyadenylation sites. The cDNA encoding QBP1(x2) was obtained from a plasmid isolated by phage display and used for protein expression in cell culture ^63^, which was fused to the cDNA encoding EGFP. The sequence of all constructs was confirmed by DNA sequencing.

Then, the QBP1(x2)-EGFP cDNA was inserted in the p-EGFP-N1 vector as a template for its production, and then it was amplified to introduce the cDNA insert into the pCAGGS vector. The entire insert with the promoter and the encoding sequence of interest was excised with SalI and PstI from the expression vector. This was performed following the similar protocols as previously described for other QBP1 constructions ^63,66^.

### Production of QBP1 transgenic mouse

Fertilized eggs derived of a C57xC57 cross were extracted from the oviduct and microinjected with the purified cDNA of the insert (enhancer-promoter-QBP1x2-EGFP-polyA terminator), by a subcontracted company (Japan SLC, Inc). A total of 200 injected ovules were implanted in pseudo-pregnant females which resulted in the birth of 3 lines of transgenic newborn mice. The incorporation of the transgene was examined by western blot in the brain lysate: brains were homogenized in 500 μL of TBS (pH 6.8) with a protease inhibitor cocktail (Nacalai Tesque). About 10 strokes using a 2 mL homogenizer was sufficient. The lysate was placed in a 2 mL tube and 600 μL of 2 x SDS sample buffer was added using a cut tip. The lysate was pipetted about 5 times, boiled for 10 min, and cooled at room temperature. 5 μL of the lysate were mixed with 20 μL of SDS sample buffer and then 10 μL were used for SDS PAGE/Western blotting.

### Recombinant protein production and amyloid characterization

The production of hCPEB3 IDR as a recombinant protein was carried out following the methodology described elsewhere ^75^. For amyloid inhibition, QBP1-M8 (octapeptide: WKWWWPGIF) and SCR-C (octapeptide: WPIWKGWF) ^64,71,72^ were prepared at 10 mM (stock concentration) in Dimethyl sulfoxide (DMSO, Sigma-Aldrich 276855) to favor its dissolution. Visualization of amyloid fibers was performed by transfer electron microscopy (TEM), where 5 µL of incubated protein (different times and concentrations) adsorbed on carbon-coated 300-mesh copper grids (Ted Pella Inc.) and subsequently were stained with 10 µL of 1% (w/v) aqueous uranyl acetate solution. The analysis was performed at 80 kV excitation voltage using a JEOL JEM 1200 EX II TEM equipped with a Megaview III CCD camera. Amyloid aggregation was monitored by performing <u>ThT</u> binding kinetics, analyzing ThT fluorescence intensity changes after binding to mature fibers ^127^. For this purpose, the reaction was prepared by mixing the sample at the desired concentration with 50 µM ThT (the stock was prepared at 1mM concentration in PBS buffer), recording fluorescence intensity measurements over time on a FLUOstar OPTIMA plate reader at 37 °C with excitation at 440 nm and emission at 490 nm. The experimental values of ThT fluorescence intensity of the samples were subtracted from the value obtained for the blank (PBS-ThT buffer). Specific details of the analysis of the homogenized fraction of seahorses are provided in the *Supplementary Information* section.

### Animals

The experiments were carried out in male and female QBP1 transgenic mice of constitutive expression from 3 to 5 months of age and wild type littermates C57BL/6Ola (WT). The mice were generated from frozen embryos donated by Prof. Yoshitaka Nagai (previously frozen by the in-house facility of National Center of Neurology and Psychiatry), starting their analysis from the third homozygous generation, whose expression of the transgene was already stable. Frozen embryos were recovered by the Mouse Embryo Cryopreservation Facility at the Centro Nacional de Biotecnología-CSIC (Madrid). The mice were housed in groups of 4-6 per cage, under conditions of constant temperature (21-22 °C) and a light/dark cycle of 12 hours; with food and water *ad libitum*. The animals have been treated following the guidelines contained in RD 53/2013 and in accordance with European Community Guidelines (Directive 2010/63/EU), previously approved by the Bioethics Committee of the Cajal Institute, The CSIC and the Madrid Community.

Genotyping was carried out by the Molecular and Cellular Biology Unit of the Cajal Institute, using mouse tail tissue (Illustra tissue and cells Genomic prep kit, GE Healthcare) and the copy number is quantified by quantitative PCR (qPCR, Taqman Copy Number assays, Applied Biosystems). A general protocol was followed with a basic battery of behavior and neurological characterization, as described elsewhere ^128^, provided by the Unit of Animal Behaviour at the Cajal Institute. Weight, temperature and abnormal physical characteristics are weekly measured (bald spots, excessive grooming and integrity of the vibrissae).

### Novel object recognition test (NOR)

NOR is a test that allows analyzing the discrimination of new objects with respect to objects already known, in a protocol of three phases: training, short-term test (both in the first day) and long-term test (second day) (N_QBP1_=9 and N_WT_=10). The protocol and the guidelines used were as described elsewhere ^129^. Mice were first habituated in the arena for 15 min and exposed to two different objects (A-B, training); 1 hour later short-term memory was tested for 10 min, in which mice are exposed to A-C objects. 24 hours later, long-term memory is tested (10 min) and mice are exposed to A-D objects. The mice were recorded using a video camera and the exploration time was counted using EthoVision XT 8.5 software: the observer who blindly analyzed the results considered the active approach of the mouse nose within a one centimeter range of the object as an exploration of the object. The discrimination index (DI) was determined by the difference in exploration time expressed as a ratio of the total time spent exploring the two objects: good discrimination is settle at 0,20 DI value, considering that there must be significant differences between the training and the testing phases of the test.

### Morris Water Maze (MWM)

After a habituation test (day 0) in which preferences between quadrants in the different experimental groups were ruled out, the animals learned to find a hidden platform during the following 5 days, through 4 trials/day (60 seconds each trial, plus 20 seconds on the platform). If an animal could not reach the platform, the experimenter placed it on top of it. Subsequently, the animals were subjected to a trial on the seventh day, without the platform, to assess preference for the platform quadrant. During this test the trajectories were analyzed to measure time spent swimming in the platform quadrant (Probe Q) compared to time in the other three quadrants, or time spent swimming above the virtual platform position (Probe P) compared to time above the virtual mirror or platform positions in the other three quadrants. On days 17 and 18, animals were exposed to a new series of four trials daily for two days, recording escape latency. To analyze the internal performance within each experimental day, we have developed a new variable called Inter-test Acquisition Ratio (IAR, **Fig. 3C**) to quantitatively compare the performance of the mice in Test 1 with respect to the average of the four tests of each acquisition day.

### Contextual Fear conditioning (CFC)

To analyze aversive memories, fear conditioning was implanted in the animals using the UGO BASILE ANY-maze controlled fear conditioning system (mouse cage). On the acquisition day, after the adaptation time (180 sec) the mice were exposed to three 30 sec trials: a light electric shock on the paw was used as unconditioned stimulus (UCS, 0.8 mA intensity) and administered in the last 2s of the trial, in which a conditioned stimulus (CS, 30s continuous white sound −70 dB-) is present. Each trial was spaced by a fixed inter-trial time (50s) and then recorded a final 30s after the last trial (Post). 24 hours after acquisition, the context test was evaluated: mice were returned to the conditioning chamber and freezing behavior was measured for 5 minutes (no shock and 250 lux). Likewise, 7 days after acquisition, the recall of the conditioned stimulus (acoustic cue) was evaluated, whose protocol is the same as the one used during fear implantation, but eliminating the unconditioned stimulus (electric shock). Extinction of fear to the conditioned context was assessed at 96 h and at 8 days after fear acquisition using the same protocol as the context test. The experimental design and subsequent data collection were performed using the behavioral tracking program associated with the equipment (ANY-maze software).

### Statistical analysis

Statistical analysis was performed with IBM SPSS Statistics (Version 26.0, Armonk, NY, IBM Corp) and GraphPad Prism (version 8.0.2). First, the normal distribution of the data was analyzed (Shapiro-Wilk, N<50), the homogeneity of their variances and the identification of extreme values, which were removed only in case they were considered extreme cases by the program analysis. Subsequently, the values were analyzed according to the number of variables and groups to be compared. In the case of comparing two samples, the Student’s t-test (unpaired), or the Mann-Whitney U-test (nonparametric cases) was used in case of independent samples; while the Student’s t-test (paired) or the Wilcoxon signed-rank test (nonparametric cases) were used in case of repeated or related measures. In case of more than one sample, a one-way ANOVA analysis of variance or the Kruskal-Wallis test (nonparametric cases) was used in case of independent samples; whereas a two-way ANOVA or mixed model (Bonferroni for posthoc) or Friedman’s test is used in case of related or paired samples. Statistical significance was set at α=0.05. The data were managed in Excel and, subsequently, graphs and images were generated using Origin 2021 and GraphPad Prism 8. For independent group comparisons: *p<0.05, **p<0.01, ***p<0.001, trends 0.05 ≥ # < 0.09; for dependent group comparisons: +p<0.05, ++p<0.01, +++p<0.001, trends 0.05 ≥ #’ < 0.09.

### Hippocampi homogenization and fractionation

In order to analyze the WT and TgQBP1 mice that have performed the hippocampal-dependent memory tasks, their hippocampi were homogenized using a detergent rich protocol ^56,58^. The extracted hippocampi were incubated in 300 μL of lysis buffer (50 mM Tris pH 7.5, 1 mM EDTA, 10 mM KCl, 0.5% Triton X-100, 0.5% NP40) for 1 hour in a cold room. Upon removing a small volume of homogenate for further analysis, the rest of the volume was centrifuged for 5 minutes at 4,000 rpm (Microfugue 18 centrifuge, Beckman Coulter) to obtain the cell debris fraction (P1) and a soluble fraction (S1), from which an aliquot was reserved too. This supernatant was ultracentrifugated at 65,000 rpm for 1 hour at 4°C in a TLA 100.1 rotor (Beckman Coulter) to obtain the soluble (S2) and insoluble (P2) protein fractions. Both pellet fractions were suspended in 50 μL of detergent buffer.

### Western blot of hippocampal samples

Dissected hippocampi were homogenized in the detergent buffer and fractionated into soluble and insoluble fractions as mentioned. From each fraction, 150 μg of protein were subjected to SDS-PAGE using 12.5% acrylamide gels and then transferred to a nitrocellulose membrane (0.45 μm, Amersham Protran) in Tris-glycine buffer with 15% methanol. PBS 0.1% Tween-20 (PBS-T) with 5% non-fat-milk (Biorad) was used to block the membrane, which was later incubated with the primary antibody: anti-CPEB3 rabbit polyclonal (ab10883) or anti-GAPDH rabbit polyclonal (ab37168). Washing steps were carried out with PBS-T 10% non-fat milk. IRDye 680LT Goat anti-rabbit (LI-COR) was used as secondary antibody and then revealed in an Odyssey® CLx (LI-COR) apparatus. The images obtained were analyzed using Image J software to quantify the intensity of the bands corresponding to the mCPEB3 band and the GAPDH loading control, for which several different Western blots were analyzed for each sample (N=5 animals/group). The analysis of these data was performed by calculating the relative intensity comparing the quantification of the mCPEB3 band with that of the GAPDH band (CPEB3/GAPDH) for each transfer and well analyzed, normalizing the quantified intensity for each band to make them comparable.

Extended information on additional experiments and detailed description of the experiments with mice has been included in the *Supplementary Information* section.

## Acknowledgements

We thank Dr. D. V. Laurents for helpful discussions and English style suggestions and to Dr. José Ignacio Robles Sánchez for critical reading of the manuscript. We thank also the Molecular and Cellular Biology Unit (UBMC) from the Cajal Institute (CSIC) for the genotyping of mice samples. P. López-García thanks the financial support of the Community of Madrid for her industrial PhD grant to DisruPep SL and CSIC (IND2020/BMD-17444). This work was supported by grants from the Spanish Ministry of Economy (SAF2013-49179-C2-1-R and SAF2016-76678-C2-1-R) to M.C.-V.

## Author Contributions

M.C.-V. conceived the project. P.L.-G, J.L.T, K.R.MG. and D.R.M. designed the animal experiments. P.L.-G, J.L.T, K.R.MG. obtained the experimental data, performed its analysis and interpreted the results with the help of A.S and A. P. Moreover, H.A.P and Y.N. designed the QBP1 construct and produced the QBP1 transgenic line, and performed the first characterization. D.R.M. recovered and maintained the initial transgenic line, checking the presence of the peptide in the TgQBP1 mice hippocampus. P.L.-G and M.E.V designed the biochemical experiments, analyzed the data and interpreted the results. Finally, P.L.-G. wrote the first draft of the manuscript, all authors read the final draft and provided feedback, and M.C.-V. wrote the final draft.

## References

1. American Psychiatric American Psychiatric Association. (2013). Diagnostic and Statistical Manual of Mental Disorders. Arlington. 10.1176/appi.books.9780890425596.744053Association. *Diagnostic and Statistical Manual of Mental Disorders*. *Arlington* (2013). doi:10.1176/appi.books.9780890425596.744053

2. National Institute of Clinical Excellence. Clinical Guideline 26: Posttraumatic stress disorder: The management of PTSD in adults and children in primary and secondary care. (2005).

3. Malikowska-Racia, N. & Salat, K. Recent advances in the neurobiology of posttraumatic stress disorder: A review of possible mechanisms underlying an effective pharmacotherapy. Pharmacol. Res. 142, 30–49 (2019).

4. Abdallah, C. G. et al. The Neurobiology and Pharmacotherapy of Posttraumatic Stress Disorder. Annu. Rev. Pharmacol. Toxicol. 59, 171–189 (2019).

5. VanElzakker, M. B. Posttraumatic Stress Disorder. Neurosci. 21st Century 4055–4084 (2016). doi:10.1007/978-1-4939-3474-4

6. Bryant, R. A. The Current Evidence for Acute Stress Disorder. Curr. Psychiatry Rep. 20, (2018).

7. Cardeña, E. & Carlson, E. Acute Stress Disorder Revisited. Annu. Rev. Clin. Psychol. 7, 245–267 (2011).

8. Kearns, M. C., Ressler, K. J., Zatzick, D. & Rothbaum, B. O. Early interventions for ptsd: a review. Depress. Anxiety 29, 833–842 (2012).

9. Atwoli, L., Stein, D. J., Koenen, K. C. & McLaughlin, K. A. Epidemiology of posttraumatic stress disorder: Prevalence, correlates and consequences. Curr. Opin. Psychiatry 28, 307–311 (2015).

10. Javidi, H. & Yadollahie, M. Post-traumatic stress disorder. Int. J. Occup. Environ. Med. 3, 2–9 (2012).

11. Yehuda, R. et al. Post-Traumatic Stress Disorder. Dis. Prim. 702–706 (2015). doi:10.1016/B978-0-08-097086-8.27051-7

12. Campos, M. R. Trastorno de estrés postraumático. 4, 233–240 (2016).

13. Howlett JR, S. M. Chapter 16. Post-Traumatic Stress Disorder: Relationship to Traumatic Brain. Chapter 16 (2016).

14. National Collaborating Centre for Mental Health. Post-Traumatic Stress Disorder: The management of PTSD in adults and children in primary and secondary care. The Royal College of Psychiatrists & The British Psychological Society 346, (2005).

15. Krystal, J. H. et al. It Is Time to Address the Crisis in the Pharmacotherapy of Posttraumatic Stress Disorder: A Consensus Statement of the PTSD Psychopharmacology Working Group. Biol. Psychiatry 82, e51–e59 (2017).

16. Koek, R. J. & Luong, T. N. Theranostic pharmacology in PTSD: Neurobiology and timing. Prog. Neuro-Psychopharmacology Biol. Psychiatry 90, 245–263 (2019).

17. Davidson, J. R. T. Pharmacologic treatment of acute and chronic stress following trauma: 2006. J. Clin. Psychiatry 67, 34–39 (2006).

18. Stein, D. J., Ipser, J. C., Seedat, S., Safer, C. & Amos, T. Pharmacotherapy for post traumatic stress disorder (PTSD). Cochrane database Syst. Rev. (2006). doi:10.1002/14651858.CD002795.pub2.www.cochranelibrary.com

19. Friedman, M. J., Donnelly, C. L. & Mellman, T. A. Pharmacotherapy for PTSD. Psychiatr Ann. 33, (2003).

20. Davidson, J. R. T. et al. Mirtazapine vs. Placebo in Posttraumatic Stress Disorder : A Pilot Trial. Biol. Psychiatry 53, 188–191 (2003).

21. Kim, H. et al. Effects of naturally occurring compounds on fibril formation and oxidative stress of beta-amyloid. J. Agric. Food Chem. 53, 8537–8541 (2005).

22. Hetrick, S. E., Purcell, R., Garner, B. & Parslow, R. Combined pharmacotherapy and psychological therapies for post traumatic stress disorder (PTSD). Cochrane Database Syst. Rev. (2010). doi:10.1002/14651858.CD007316.pub2

23. Coventry, P. A., et al. Psychological and pharmacological interventions for posttraumatic stress disorder and comorbid mental health problems following complex traumatic events: Systematic review and component network meta-analysis. PLoS Medicine 17, (2020).

24. Lee, D. J. et al. Psychotherapy Versus Pharmacotherapy for Posttraumatic Stress Disorder: Systemic Review and Meta-Analyses To Determine First-Line Treatments. Uniformed Serv. Univ. Heal. Sci. 166, (2016).

25. Astill Wright, L., et al. Pharmacological prevention and early treatment of post-traumatic stress disorder and acute stress disorder: a systematic review and meta-analysis. Transl. Psychiatry 9, 1–10 (2019).

26. Koek, R. J. & Luong, T. N. Theranostic pharmacology in PTSD: Neurobiology and timing. Prog. Neuro-Psychopharmacology Biol. Psychiatry 90, 245–263 (2019).

27. Kindt, M. & van Emmerik, A. New avenues for treating emotional memory disorders: towards a reconsolidation intervention for posttraumatic stress disorder. Ther. Adv. Psychopharmacol. 6, 283–295 (2016).

28. Yan, Y. et al. Neuronal Circuits Associated with Fear Memory: Potential Therapeutic Targets for Posttraumatic Stress Disorder. Neuroscientist 29, 332–351 (2023).

29. Maddox, S. A., Hartmann, J., Ross, R. A. & Ressler, K. J. Deconstructing the Gestalt: Mechanisms of Fear, Threat, and Trauma Memory Encoding. Neuron 102, 60–74 (2019).

30. Villain, H., Benkahoul, A., Birmes, P., Ferry, B. & Roullet, P. Influence of early stress on memory reconsolidation: Implications for posttraumatic stress disorder treatment. PLoS One 13, (2018).

31. Visser, R. M., Lau-Zhu, A., Henson, R. N. & Holmes, E. A. Multiple memory systems, multiple time points: how science can inform treatment to control the expression of unwanted emotional memories. Phil. Trans. R. Soc. B 373, 20170209 (2018).

32. Careaga, M. B. L., Girardi, C. E. N. & Suchecki, D. Understanding posttraumatic stress disorder through fear conditioning, extinction and reconsolidation. Neurosci. Biobehav. Rev. 71, 48–57 (2016).

33. Cohen, H. et al. Traumatic Memory Consolidation and Attenuates Posttraumatic Stress Response in Rats. Biol. Psychiatry 60, 767–776 (2006).

34. Wang, X. et al. Transcriptional Regulation Involved in Fear Memory Reconsolidation. J. Mol. Neurosci. (2018).

35. Torregrossa, M. M. & Taylor, J. R. Learning to forget: Manipulating extinction and reconsolidation processes to treat addiction. Psychopharmacology (Berl). 226, 659–672 (2013).

36. Aliev, G. et al. Neurophysiology and Psychopathology Underlying PTSD and Recent Insights into the PTSD Therapies—A Comprehensive Review. J. Clin. Med. 9, 2951 (2020).

37. Sijbrandij, M., Kleiboer, A., Bisson, J. I., Barbui, C. & Cuijpers, P. Pharmacological prevention of post-traumatic stress disorder and acute stress disorder: A systematic review and meta-analysis. The Lancet Psychiatry 2, 413–421 (2015).

38. Monk, C. S. & Nelson, C. A. The Effects of Hydrocortisone on Cognitive and Neural Function : A Behavioral and Event-Related Potential Investigation. Neuropsychopharmacology 26, 505–519 (2002).

39. Olff, M. et al. Social support, oxytocin, and PTSD. Eur. J. Psychotraumatol. 5, 26513 (2014).

40. Kirsch, P. et al. Oxytocin modulates neural circuitry for social cognition and fear in humans. J. Neurosci. 25, 11489–11493 (2005).

41. Yoon, S. & Kim, Y. K. Neuroendocrinological treatment targets for posttraumatic stress disorder. Prog. Neuro-Psychopharmacology Biol. Psychiatry 90, 212–222 (2019).

42. Kindt, M. & van Emmerik, A. New avenues for treating emotional memory disorders: towards a reconsolidation intervention for posttraumatic stress disorder. Ther. Adv. Psychopharmacol. 6, 283–295 (2016).

43. Wood, N. E. et al. Pharmacological blockade of memory reconsolidation in posttraumatic stress disorder: Three negative psychophysiological studies. Psychiatry Res. 225, 31–39 (2015).

44. Giustino, T. F., Fitzgerald, P. J. & Maren, S. Revisiting propranolol and PTSD: Memory erasure or extinction enhancement? Neurobiol Learn Mem 130, 26–33 (2017).

45. Pitman, R. K. et al. Pilot Study of Secondary Prevention of Posttraumatic Stress Disorder with Propranolol. Biol. Psychiatry 51, 189–142 (2002).

46. Birk, J. L. et al. Early interventions to prevent posttraumatic stress disorder symptoms in survivors of life-threatening medical events: A systematic review. J. Anxiety Disord. 64, 24–39 (2019).

47. Si, K. & Kandel, E. R. The role of functional prion-like proteins in the persistence of memory. Cold Spring Harb. Perspect. Biol. 8, (2016).

48. Li, L. et al. A Putative Biochemical Engram of Long-Term Memory. Curr. Biol. 26, 3143–3156 (2016).

49. Nizhnikov, A. A., Antonets, K. S., Bondarev, S. A., Inge-Vechtomov, S. G. & Derkatch, I. L. Prions, amyloids, and RNA: Pieces of a puzzle. Prion 10, 182–206 (2016).

50. Lu, W. H. et al. CPEB3-dowregulated Nr3c1 mRNA translation confers resilience to developing posttraumatic stress disorder-like behavior in fear-conditioned mice. Neuropsychopharmacology 1–11 (2021). doi:10.1038/s41386-021-01017-2

51. Theis, M., Si, K. & Kandel, E. R. Two previously undescribed members of the mouse CPEB family of genes and their inducible expression in the principal cell layers of the hippocampus. Proc. Natl. Acad. Sci. U. S. A. 100, 9602–9607 (2003).

52. Richter, J. D. CPEB: a life in translation. Trends Biochem. Sci. 32, 279–285 (2007).

53. Mendez, R. & Richter, J. D. Translational control by CPEB: A means to the end. Nat. Rev. Mol. Cell Biol. 2, 521–529 (2001).

54. Chen, M., Zheng, W. & Wolynes, P. G. Energy landscapes of a mechanical prion and their implications for the molecular mechanism of long-term memory. Proc. Natl. Acad. Sci. 113, 5006–5011 (2016).

55. Krüttner, S. et al. Synaptic Orb2A Bridges Memory Acquisition and Late Memory Consolidation in Drosophila. Cell Rep. 11, 1953–1965 (2015).

56. Stephan, J. S. et al. The CPEB3 protein is a Functional prion that interacts with the actin cytoskeleton. Cell Rep. 11, 1772–1785 (2015).

57. Raveendra, B. L. et al. Characterization of prion-like conformational changes of the neuronal isoform of Aplysia CPEB. Nat. Struct. Mol. Biol. 20, 495–501 (2013).

58. Drisaldi, B. et al. SUMOylation is an inhibitory constraint that regulates the prion-like aggregation and activity of CPEB3. Cell Rep. 11, 1694–1702 (2015).

59. Pavlopoulos, E. et al. Neuralized1 activates CPEB3: A function for nonproteolytic ubiquitin in synaptic plasticity and memory storage. Cell 147, 1369–1383 (2011).

60. Fioriti, L. et al. The persistence of hippocampal-based memory requires protein synthesis mediated by the prion-like protein CPEB3. Neuron 86, 1433–1448 (2015).

61. Tsai, L. Y. et al. CPEB4 knockout mice exhibit normal hippocampus-related synaptic plasticity and memory. PLoS One 8, (2013).

62. Lu, W. H., Yeh, N. H. & Huang, Y. S. CPEB2 Activates GRASP1 mRNA Translation and Promotes AMPA Receptor Surface Expression, Long-Term Potentiation, and Memory. Cell Rep. 21, 1783–1794 (2017).

63. Nagai, Y. et al. Inhibition of polyglutamine protein aggregation and cell death by novel peptides identified by phage display screening. J. Biol. Chem. 275, 10437–10442 (2000).

64. Popiel, H. A., Nagai, Y., Fujikake, N. & Toda, T. Protein transduction domain-mediated delivery of QBP1 suppresses polyglutamine-induced neurodegeneration in vivo. Mol. Ther. 15, 303–309 (2007).

65. Popiel, H. A. et al. The Aggregation Inhibitor Peptide QBP1 as a Therapeutic Molecule for the Polyglutamine Neurodegenerative Diseases. J. Amino Acids 2011, 1–10 (2011).

66. Popiel, H. A. et al. Inhibition of Protein Misfolding/Aggregation Using Polyglutamine Binding Peptide QBP1 as a Therapy for the Polyglutamine Diseases. Neurotherapeutics 10, 440–446 (2013).

67. Nagai, Y. et al. A toxic monomeric conformer of the polyglutamine protein. Nat. Struct. Mol. Biol. 14, 332–340 (2007).

68. Nagai, Y. et al. Prevention of polyglutamine oligomerization and neurodegeneration by the peptide inhibitor QBP1 in Drosophila. Hum. Mol. Genet. 12, 1253–1260 (2003).

69. Hervás, R. et al. Common features at the start of the neurodegeneration cascade. PLoS Biol. 10, (2012).

70. Mompeán, M., Ramírez de Mingo, D. & Hervás, R. Molecular mechanism of the inhibition of TDP-43 amyloidogenesis by QBP1. Arch. Biochem. Biophys. (2019).

71. Tomita, K. et al. Structure-activity relationship study on polyglutamine binding peptide QBP1. Bioorganic Med. Chem. 17, 1259–1263 (2009).

72. Ramos-Martín, F., Hervás, R., Carrión-Vázquez, M. & Laurents, D. V. NMR spectroscopy reveals a preferred conformation with a defined hydrophobic cluster for polyglutamine binding peptide 1. Arch. Biochem. Biophys. 558, 104–110 (2014).

73. Hervás, R. et al. Divergent CPEB prion-like domains reveal different assembly mechanisms for a generic amyloid-like fold. (2020). doi:10.1101/2020.05.19.103804

74. Hervás, R. et al. Molecular basis of Orb2 amyloidogenesis and blockade of memory consolidation. PLoS Biol. 14, 1–32 (2016).

75. Ramírez de Mingo, D., et al. Phase separation modulates the functional amyloid assembly of human CPEB3. Prog. Neurobiol. 231, (2023).

76. Kayed, R. et al. Fibril specific, conformation dependent antibodies recognize a generic epitope common to amyloid fibrils and fibrillar oligomers that is absent in prefibrillar oligomers. Mol. Neurodegener. 2, 1–11 (2007).

77. Okamoto, Y. et al. Surface plasmon resonance characterization of specific binding of polyglutamine aggregation inhibitors to the expanded polyglutamine stretch. Biochem. Biophys. Res. Commun. 378, 634–639 (2009).

78. Antunes, M. & Biala, G. The novel object recognition memory: Neurobiology, test procedure, and its modifications. Cogn. Process. 13, 93–110 (2012).

79. Aliczki, M. & Haller, J. Electric Shock as Model of Post-traumatic Stress Disorder in Rodents. In: Martin C., Preedy V., Patel V. (eds) Comprehensive Guide to Post-Traumatic Stress Disorder. Springer, Cham. 1–16 (2015). doi:10.1007/978-3-319-08613-2

80. Lee, F. S. et al. Adolescent mental health--Opportunity and obligation. Science (80-.). 346, 547–549 (2014).

81. Pattwell, S. S., Casey, B. J. & Lee, F. S. The Teenage Brain: Altered Fear in Humans and Mice. Curr. Dir. Psychol. Sci. 22, 146–151 (2013).

82. Bisby, M. A., Stylianakis, A. A., Baker, K. D. & Richardson, R. Fear extinction learning and retention during adolescence in rats and mice: A systematic review. Neurosci. Biobehav. Rev. 131, 1264–1274 (2021).

83. Bauer, E. P. Sex differences in fear responses: Neural circuits. Neuropharmacology 222, 109298 (2023).

84. Gradus, J. L. Prevalence and prognosis of stress disorders: A review of the epidemiologic literature. Clin. Epidemiol. 9, 251–260 (2017).

85. Russo, A. S. & Parsons, R. G. Behavioral Expression of Contextual Fear in Male and Female Rats. Front. Behav. Neurosci. 15, 1–9 (2021).

86. Keiser, A. A. et al. Sex Differences in Context Fear Generalization and Recruitment of Hippocampus and Amygdala during Retrieval. Neuropsychopharmacology 42, 397–407 (2017).

87. Lynch, J., Cullen, P. K., Jasnow, A. M. & Riccio, D. C. Sex differences in the generalization of fear as a function of retention intervals. Learn. Mem. 20, 628–632 (2013).

88. Klunk, W. E., Pettergrew, J. W. & Abraham, D. J. Quatitative Evaluation of Congo Red Binding with a Beta-pleated to. J. Histochem. Cytochem. 37, 1273–1281 (1989).

89. Linke, R. P. Congo red staining of amyloid: improvements and practical guide for a more precise diagnosis of amyloid and the different amyloidoses. Protein Misfolding, Aggregation, Conform. Dis. 239–276 (2006). doi:10.1007/b136464

90. Chao, H.-W. et al. Deletion of CPEB3 Enhances Hippocampus-Dependent Memory via Increasing Expressions of PSD95 and NMDA Receptors. J. Neurosci. 33, 17008–17022 (2013).

91. Ford, L., Ling, E., Kandel, E. R. & Fioriti, L. CPEB3 inhibits translation of mRNA targets by localizing them to P bodies. PNAS 116, 18078–18087 (2019).

92. Vermetten, E. & Bremner, J. D. Circuits and systems in stress. II. Applications to neurobiology and treatment in posttraumatic stress disorder. Depress. Anxiety 16, 14–38 (2002).

93. Abdallah, C. G. et al. The neurobiology and pharmacotherapy of posttraumatic stress disorder. Annu. Rev. Pharmacol. Toxicol. 59, 171–189 (2019).

94. Gu, W., Wang, C., Li, Z., Wang, Z. & Zhang, X. Pharmacotherapies for posttraumatic stress disorder a meta-analysis. J. Nerv. Ment. Dis. 204, 331–338 (2016).

95. Nagai, Y. et al. Inhibition of polyglutamine protein aggregation and cell death by novel peptides identified by phage display screening. J. Biol. Chem. 275, 10437–10442 (2000).

96. Takeuchi, T. & Nagai, Y. Protein misfolding and aggregation as a therapeutic target for polyglutamine diseases. Brain Sci. 7, (2017).

97. Flores, M. D. et al. Structure of a reversible amyloid fibril formed by the CPEB3 prion-like domain reveals a core sequence involved in translational regulation. bioRxiv Prepr. (2022).

98. Reselammal, D. S. et al. Mapping the fibril core of the prion subdomain of the mammalian CPEB3 that is involved in long term memory retention. J. Mol. Biol. 433, 167084 (2021).

99. Ramírez de Mingo, D., Pantoja-Uceda, D., Hervás, R., Carrión-Vázquez, M. & Laurents, D. V. Conformational dynamics in the disordered region of human CPEB3 linked to memory consolidation. BMC Biol. 20, 1–18 (2022).

100. Belwal, V. K., Datta, D. & Chaudhary, N. The β-turn-supporting motif in the polyglutamine binding peptide QBP1 is essential for inhibiting huntingtin aggregation. FEBS Lett. 594, 2894–2903 (2020).

101. Hervás, R. et al. Divergent CPEB prion-like domains reveal different assembly mechanisms for a generic amyloid-like fold. BMC Biol. 19, 1–14 (2021).

102. Kamagata, K. et al. Suppression of TDP-43 aggregation by artificial peptide binder targeting to its low complexity domain. Biochem. Biophys. Res. Commun. 662, 119–125 (2023).

103. Nagai, Y. et al. Prevention of polyglutamine oligomerization and neurodegeneration by the peptide inhibitor QBP1 in Drosophila. Hum. Mol. Genet. 12, 1253–1260 (2003).

104. Zhan, L., Guo, D., Chen, G. & Yang, J. Effects of repetition learning on associative recognition over time: Role of the hippocampus and prefrontal cortex. Front. Hum. Neurosci. 12, 1–14 (2018).

105. Sanguanini, M. & Cattaneo, A. A continuous model of physiological prion aggregation suggests a role for Orb2 in gating long-term synaptic information. R. Soc. Open Sci. 5, (2018).

106. Li, L., McGinnis, J. P. & Si, K. Translational Control by Prion-like Proteins. Trends Cell Biol. 28, 494–505 (2018).

107. Myers, K. M., Ressler, K. J. & Davis, M. Different mechanisms of fear extinction dependent on length of time since fear acquisition. Learn. Mem. 13, 216–223 (2006).

108. Gradari, S. et al. The relationship between behavior acquisition and persistence abilities: Involvement of adult hippocampal neurogenesis. Hippocampus 26, 857–874 (2016).

109. Qu, W. et al. CPEB3 regulates neuron-specific alternative splicing and involves neurogenesis gene expression. Aging (Albany. NY). 13, 2330–2347 (2021).

110. Wang, X. P. & Cooper, N. G. F. Characterization of the transcripts and protein isoforms for cytoplasmic polyadenylation element binding protein-3 (CPEB3) in the mouse retina. BMC Mol. Biol. 10, 1–19 (2009).

111. Baran, S. E., Armstrong, C. E., Niren, D. C., Hanna, J. J. & Conrad, C. D. Chronic stress and sex differences on the recall of fear conditioning and extinction. Neurobiol. Learn. Mem. 91, 323–332 (2009).

112. Fleischer, A. W. & Frick, K. M. New perspectives on sex differences in learning and memory. Trends Endocrinol. Metab. 34, 526–538 (2023).

113. Baltaci, S. B., Mogulkoc, R. & Baltaci, A. K. Molecular Mechanisms of Early and Late LTP. Neurochem. Res. 44, 281–296 (2019).

114. Kitamura, T. et al. Engrams and circuits crucial for systems consolidation of a memory. Science (80-.). 356, 73–78 (2017).

115. Kandel, E. R. The molecular biology of memory storage: a dialog between genes and synapses. Biosci. Rep. 24, 475–522 (2005).

116. Squire, L. R. Lost forever or temporarily misplaced? The long debate about the nature of memory impairment. Learn. Mem. 13, 522–529 (2006).

117. Lynch, G., Kramár, E. A. & Gall, C. M. Protein synthesis and consolidation of memory-related synaptic changes. Brain Res. 1621, 62–72 (2015).

118. Huang, W. H., Chao, H. W., Tsai, L. Y., Chung, M. H. & Huang, Y. S. Elevated activation of CaMKIIα in the CPEB3-knockout hippocampus impairs a specific form of NMDAR-dependent synaptic depotentiation. Front. Cell. Neurosci. 8, 1–12 (2014).

119. Morrison, F. G. & Ressler, K. J. From the neurobiology of extinction to improved clinical treatments. Depress. Anxiety 31, 279–290 (2014).

120. Haider, S., Batool, Z. & Rafiq, S. Method for the identification of pharmacological intervention for the disruption of fear memory in PTSD-rat model. MethodsX 7, 101059 (2020).

121. Giustino, T. F., Fitzgerald, P. J. & Maren, S. Revisiting propranolol and PTSD: Memory erasure or extinction enhancement? Neurobiol Learn Mem 130, 26–33 (2016).

122. Lonergan, M. H., Olivera-Figueroa, L. A., Pitman, R. K. & Brunet, A. Propranolol’s effects on the consolidation and reconsolidation of long-term emotional memory in healthy participants: A meta-analysis. J. Psychiatry Neurosci. 38, 222–231 (2013).

123. Brunet, A. et al. Reduction of PTSD Symptoms With Pre-Reactivation Propranolol Therapy : A Randomized Controlled Trial. Am. J. Psychiatry 175, 427–433 (2018).

124. Careaga, M. B. L., Girardi, C. E. N. & Suchecki, D. Propranolol failed to prevent severe stress-induced long-term behavioral changes in male rats. Prog. Neuro-Psychopharmacology Biol. Psychiatry 110079 (2020). doi:10.1016/j.pnpbp.2020.110079

125. Amos, T., Stein, D. & Ipser, J. Pharmacological interventions for preventing post-traumatic stress disorder (PTSD) (Review). Cochrane database Syst. Rev. CD006239 (2014). doi:10.1002/14651858.CD006239.pub2.www.cochranelibrary.com

126. Bowers, M. E. & Ressler, K. J. An overview of translationally informed treatments for PTSD: animal models of Pavlovian fear conditioning to human clinical trials. Physiol. Behav. 78, 100–106 (2015).

127. Sabate, R., Rodriguez-Santiago, L., Sodupe, M., Saupe, S. J. & Ventura, S. Thioflavin-T excimer formation upon interaction with amyloid fibers. Chem. Commun. 49, 5745 (2013).

128. Crawley, J. N. Behavioral phenotyping of transgenic and knockout mice: Experimental design and evaluation of general health, sensory functions, motor abilities, and specific behavioral tests. Brain Res. 835, 18–26 (1999).

129. McGreevy, K. R. et al. Intergenerational transmission of the positive effects of physical exercise on brain and cognition. Proc. Natl. Acad. Sci. 116, 201816781 (2019).

