## Supplementary Information for "An anti-amyloidogenic approach to specifically block memory consolidation in mice for therapeutic intervention"

##### *Supplementary experimental data*

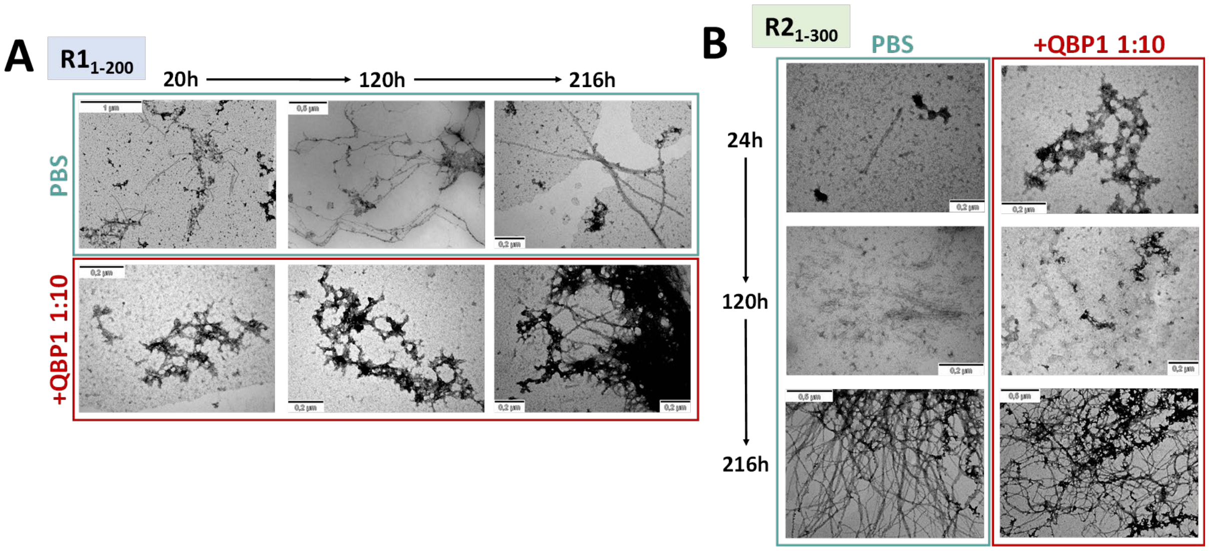


**Figure S1. QBP1-M8 induces non-specific aggregation during hCPEB3 fibrillation.**
**A)** Representative TEM micrographs of the amyloid cascade of hCPEB3 $R1_{1\text{-}200}$ (10 µM) in the presence or absence of QBP1-M8 (1:10). Samples incubated at 37 ºC in PBS pH 7.4 without shaking. The scale bar is indicated on each micrograph.
**B)** Representative images of fibrillation of de hCPEB3 $R2_{1\text{-}300}$ (10 µM) in the presence or absence of QBP1-M8 (1:10). Samples incubated at 37 ºC in PBS pH 7.4 without shaking. Scale bar is indicated on each micrograph.

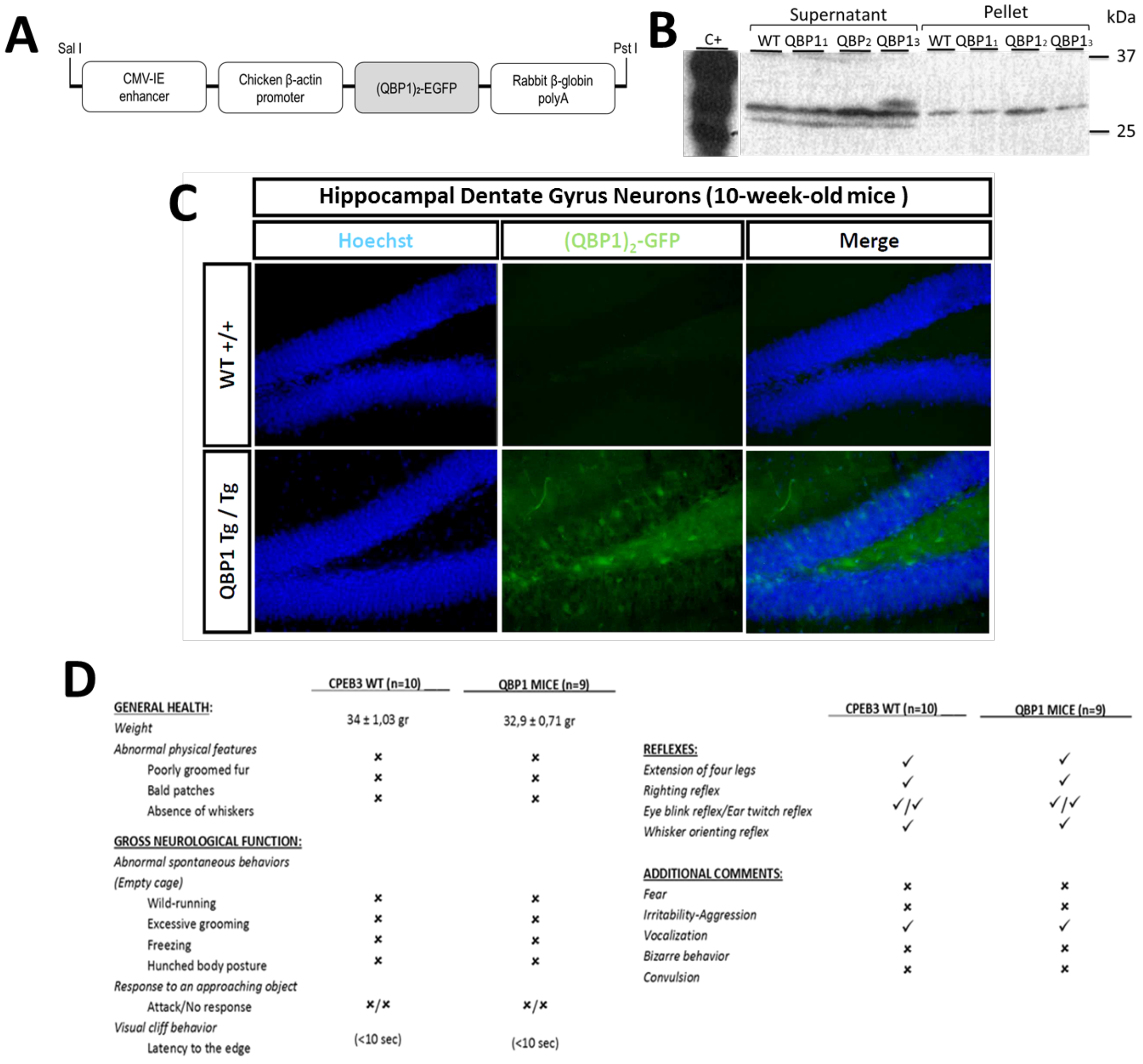


**Figure S2: QBP1 transgenic mouse: generation, characterization and QBP1 expression.**

**A)** Schematic representation of the construct used to generate the QBP1 transgenic mouse. This construct was used to microinject fertile eggs into pseudo-pregnant females, using the CAG promoter for its constitutive expression and long-term maintenance of protein expression.

**B)** Western blot of the generated transgenic lines. Of the three lines produced, only one of them (named QBP1-3) confirmed the expression of the peptide in the supernatant and was the one used to generate the breeding mouse. Q19-YFP expressing COS-7 lysate was used as a positive control, revealed with an anti-GFP (1:500) and a secondary mouse IgG-HRP antibody (1:2000).

**C)** Confocal microscopy images of hippocampal dentate gyrus sections obtained from WT and TgQBP1 mice (10 weeks old), in which EGFP was observed for its native fluorescence and Hoechst, as a DNA stain, was used to label cell nuclei. Scale bar: 50 µm.

**D)** General health characterization of the QBP1 transgenic mouse according to Crawley, 1999[1].

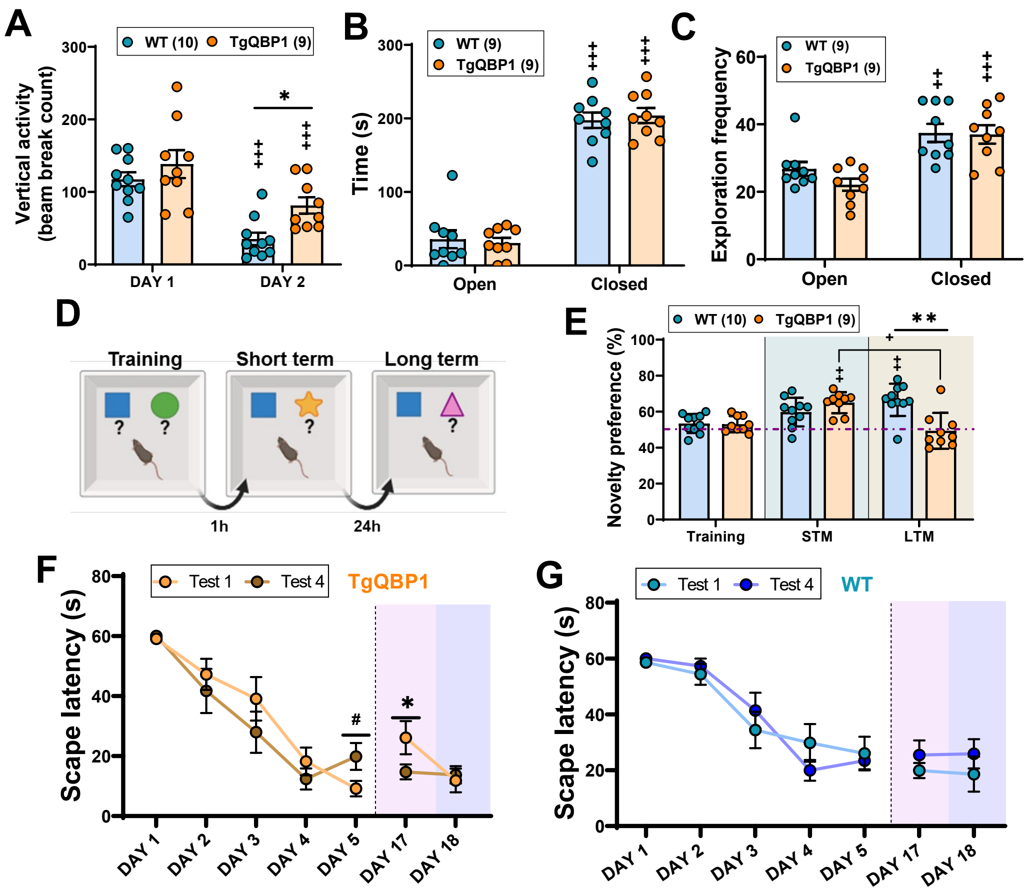


**Figure S3. Extended characterization of long-term memory in the TgQBP1 mouse.**

**A)** Total vertical activity on the two days of the protocol in the activity cage, where on day 1 there was no difference between groups. On re-exposure on day 2, the TgQBP1 mouse significantly increased its activity compared to the WT mouse (p=0.027), despite both significantly decreasing their activity compared to the previous day (WT p<0.001 and TgQBP1 p<0.001).

**B)** Time spent in the open and closed arms of the elevated cross maze in basal conditions, where TgQBP1 mice show normal anxiety behavior. There is no significant difference between groups in total time, where both spent significantly more time in the closed arms than in the open arms (p<0.001).

**C)** Exploration in the elevated cross maze in basal conditions, where the mobility of the animals is evident and they enter significantly more times into the closed arms (TgQBP1 p<0.001 and WT p=0.0012).

**D)** Object Recognition Protocol overview: The test involves a training phase (15 min) where a mouse freely explores a box with two distinct objects. After one hour (short term), the mouse explores the box again (10 min) with one familiar (blue) and one new (yellow) object. Twenty-four hours later (long term), the mouse is reintroduced into the box with the same familiar object (blue) and a new one (pink). The mouse must discriminate the novel object by increased exploration.

**E)** The percentage of preference for novelty reflects that the mice modify their preference as a function of time and group (Interaction p<0.001). The WT mouse increased its preference from training to long term (p=0.002), while the TgQBP1 mouse significantly increases its preference from training to short term (p=0.004) and this significantly decreases from short to long term (p=0.041). This is because in the long term, the WT mouse prefers the novelty significantly more than the TgQBP1 mouse (p=0.003), because it divides its preference between both objects (50-50%). The dash purple line indicates the neutral value of preference (50%), from which novelty is discriminated.

**F)** Performance plot of TgQBP1 mice in test acquisition: the escape latency corresponding to the first trial of each day (Test 1) and the last trial of each day (Test 4) are shown. These data show that the TgQBP1 mouse performs better on each test even within the same day, as evidenced by a significant trend on Day 5 (p=0.063) and a significant difference on Day 17 (p=0.040). Data were analyzed with the Mann Whitney test.

**G)** Performance plot of WT mice in test acquisition: the escape latency corresponding to the first trial of each day (Test 1) with respect to the last test of each day (Test 4) is shown. Data show that the WT mouse maintains a similar performance in all the tests of each day, with no significant differences between them.

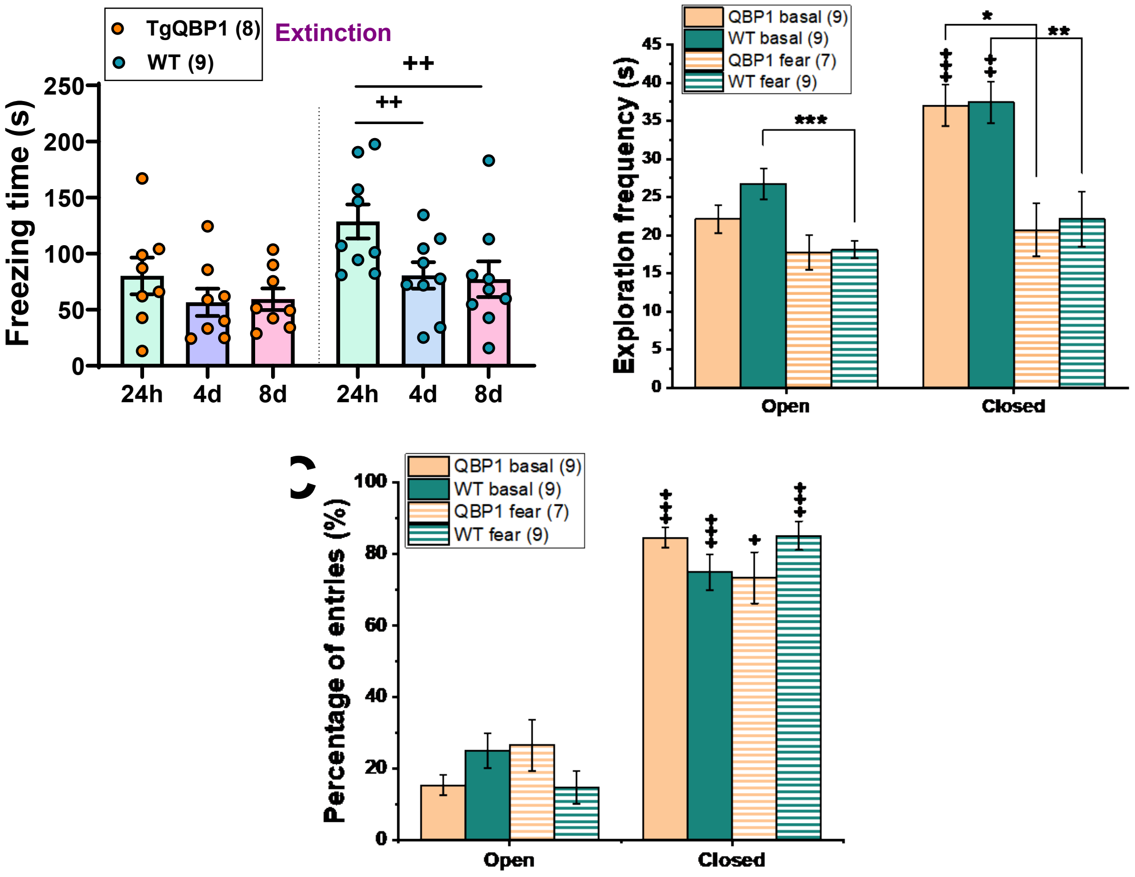


**Figure S4: Contextual fear conditioning confirmed the blockade of aversive memories in TgQBP1 mice.**
**A)** Comparison of the total time of fear expressed by the animals in the extinction of the conditioning to the aversive context: at 24h, 4 days and 8 days after acquisition. At a global level, the animals extinguished the aversive memory ($p<0.001$), although only the WT mouse significantly reduced its fear time in the successive exposures with respect to 24h (4d 0=0.006 and 8d p=0.009).
**B)** Frequency of exploration in the raised cross maze after aversive conditioning. These results show that both groups significantly decreased their explorations in the closed arms after aversive conditioning ($p=0.005$ and $p=0.016$, respectively) and only the WT mice decreased theirs in open arms ($p=0.001$), evidencing lower exploration due to higher anxiety after conditioning. Intra-subject differences showed that both groups significantly increased their exploration in the closed arms under basal conditions ($p<0.001$ TgQBP1 and $p=0.008$ WT).
**C)** Percentage of entries in the raised cross maze arms after aversive conditioning. There were no differences between groups, although within each group their percentage of entries in closed arms was significantly increased for both conditions (TgQBP1: $p<0.001$ basal and $p=0.028$ aversive; WT: $p= 0.001$ basal and $p<0.001$ aversive).
For comparisons between independent groups: *p<0.05, **p<0.01, ***p<0.001, trends 0.05 ≥ #p<0.09; for comparisons between dependent groups: +p<0.05, ++p<0.01, +++p<0.001, trends 0.05 ≥ #'p < 0.09.

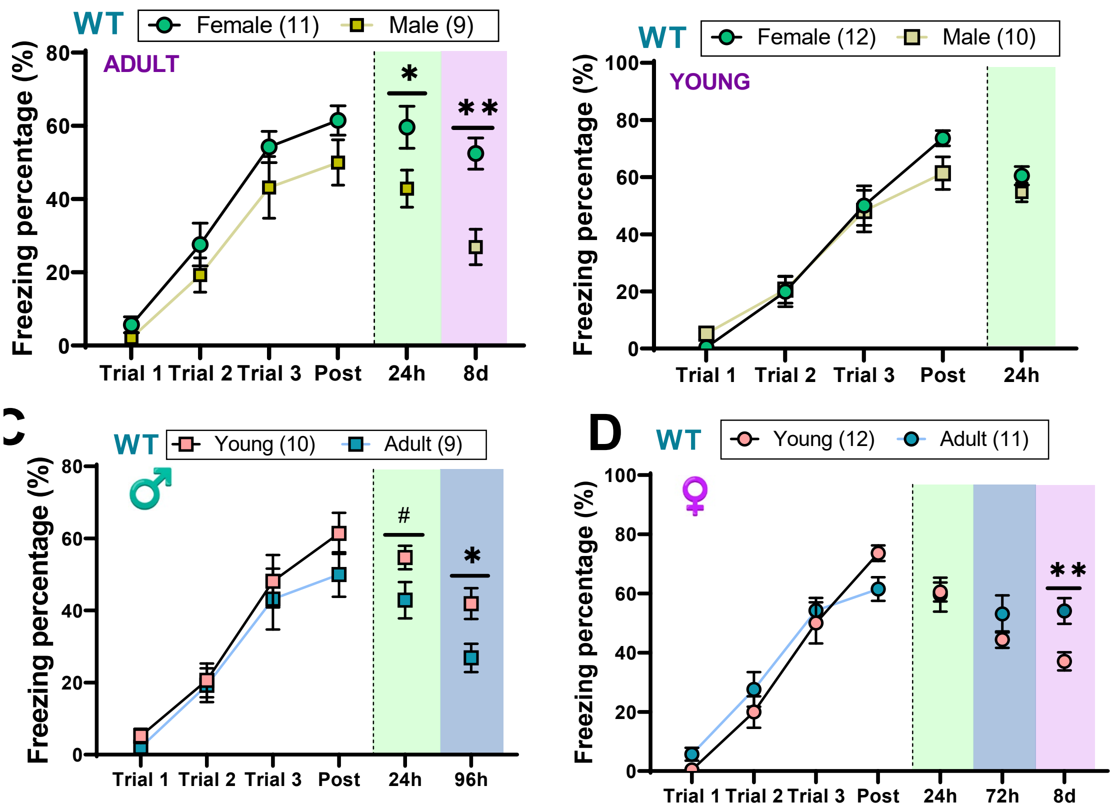


**Figure S5. Effect of age and sex of WT mice on fear conditioning.**

**A)** Effect of gender in the adult WT mouse on aversive conditioning. In the acquisition phase, the two groups significantly learn conditioning with trials (T) and there were no differences between them. However, the percentage of total fear in the review of the aversive context at 24 h reflected that adult males remember significantly worse the fear with respect to adult females (p=0.045), which is maintained in the review of the context at 8 days (p=0.009).

**B)** Effect of sex in the young WT mouse on aversive conditioning. In the acquisition phase, the two groups significantly learned conditioning with trials (T) and there are no differences between them. The absence of differences was maintained in the context review at 24 h, reflecting that there is no sex difference in the young animals.

**C)** Effect of age on the consolidation of aversive memories in WT mouse males, where both groups significantly learned the conditioning with acquisition (p<0.001. However, in the review of context fear at 24 h and 96 h the young WT mouse showed significantly more fear than the adult (p=0.061 and p=0.021 respectively).

**D)** Effect of age on the consolidation of aversive memories in females of the WT mouse, showing that during acquisition both groups significantly learned the conditioning (p<0.001) and there were no significant differences between age groups in the review of the context at 24 h and 72 h. However, adult female WT mice showed more freezing than young female mice in the context review at 8 d post acquisition (0,004).

For comparisons between independent groups: *p<0.05, **p<0.01, ***p<0.001, trends 0.05 ≥ # < 0.09; for comparisons between dependent groups: +p<0.05, ++p<0.01, +++p<0.001, trends 0.05 ≥ #' < 0.09.

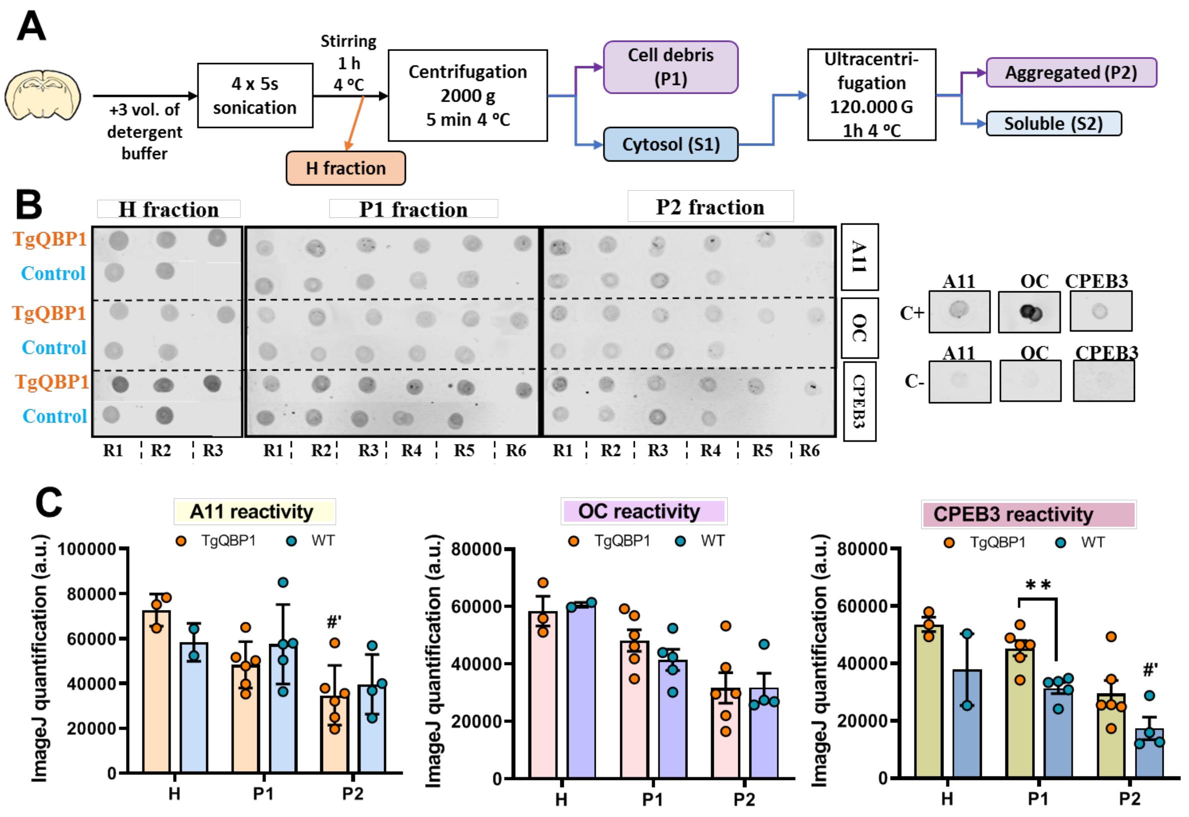


**Figure S6: Additional biochemical studies showing differences among fractions and conditions.**

**A)** Scheme of the strong extraction protocol for the homogenization of mouse hippocampi. After adding the buffer with detergents, the material is sonicated for further extraction. After one hour of shaking on ice, a first gentle centrifugation of the homogenate is performed, obtaining a soluble fraction (cytosolic fraction, S1) and an insoluble fraction (cell membrane fraction, S2). This supernatant (cytosol) is ultracentrifuged in cold to obtain the fractions corresponding to the aggregated proteins (precipitate, P2) and soluble proteins (supernatant, S2).

**B) I**mmunodot blot of the insoluble fractions of the extracted hippocampi after learning, revealed with the A11/OC conformational antibodies and with CPEB3 antibody as a loading control. The samples analyzed are listed at the bottom: R refers to mouse and the number refers to the number of samples analyzed for this group. Aβ and BSA were used as a positive and negative a control, respectively.

**C)** Quantification of immunodot blot samples with Image J software for each primary antibody used. The statistical analysis shows that there are no differences between groups in reactivity against A11/OC conformational antibodies, although there is a significant difference in the immunoreactivity with CPEB3 in the P1 fraction ($p=0.007$), where there is significantly more protein in the samples from the TgQBP1 mouse. The quantification of Aβ and BSA was used as positive and negative control, respectively.

### *Extended materials and methods*

**Analysis of hippocampal samples by Thioflavin-T fluorescence intensity (ThT)**
The assay was performed on Corning-384 plates, maintaining a final 20 µL volume: 1/6 dilution of the hippocampal extracts was analyzed. A continuous measurement was performed during 7 hours, programming the measurements every 120 seconds with a gain per well of 1300 at the FLUOStar OPTIMA Microplate Reader (BMG LABTECH), using two filters: a 440nm for excitation and 490 nm for emission. Buffer with the same amount of ThT was used as a control for fluorescence background.

**Immunodot blot assay**
This technique was used for the detection of the amyloid conformation and also the identification of the CPEB3 protein in their native state immobilized in a nitrocellulose membrane. We have analyzed their binding to antibodies that recognize specific structural conformations [2]: A11 antibody, which detects the existence of prefibrillar (toxic) oligomers, and OC antibody, which detects fibrillar oligomers and mature fibers. Also we have used an anti-CPEB3 antibody specific for the sequence of this protein (Abcam, ab10883). Extracted hippocampal samples where diluted 1/6 and then 2 µl were placed on a 0.45 µm nitrocellulose membrane (Amersham Protran). The membrane was blocked using 10% fat-free milk (Blotting-Grade Blocker non-fat dry milk, Bio-Rad) and subsequently incubated with the corresponding primary antibody: A11 antibody diluted 1:2000 (Invitrogen, ref: AHB0052), OC antibody diluted 1:2500 (Millipore, AB2286) and CPEB3 antibody diluted 1:1000 (Abcam, ab10883). A fluorescently labeled anti-rabbit secondary antibody (IRDye 680RD, Li-Cor) was used for development on an Odyssey® CLx apparatus (LI-COR). Pre-fibrillar oligomers and fibrillar species of Aβ42 was used as positive control.

**Semi-denaturing detergent agarose gel electrophoresis (SDD-AGE)**
Samples of 50 µM hCEPB3 IDR (variable final volume, 50-100 µL) incubated at different times and in the presence/absence of different inhibitors were mixed with DNA loading buffer and 2% SDS was added. A total of 10-15 µL of the mixture was loaded onto 1.5% agarose gels (TAE 1x 0.1% SDS) and the electrophoresis was performed at 70 V for 2 h at 4 °C. Proteins were then transferred to a nitrocellulose membrane filter using a Trans-Blot device (Bio-Rad) and incubated with anti-OC (1:2500, Millipore) and anti-His tag (1:2000, Novagen) antibodies. Finally, the membrane was incubated with the LI-COR antibodies as described for the western blots and dot blots.

**Activity cage and Open Field (OF)**
The spontaneous motor activity of the animals was measured with an actimeter (VersaMax Legacy Open Field activity box, Omnitech Electronics), connected to a VersaMax analyzer and the VersaDat software. Two mice were placed *per* activity cage and the measurement was set up in five cycles of 1 minute each, in a two-day protocol. Animals were habituated to the room for 30 minutes before the experiment and the cage was cleaned with ethanol before and between animals. Same protocol was used for Open field test, using one mice *per* cage and it was set up in two cycles of 5 minutes each, done in a single day.

**Elevated plus maze (EPM)**
The testing apparatus consists of four black polypropylene arms (Coulbourn Instruments, Whitehall, PA), with two "open" arms and two "closed" arms (30 cm high walls); all 10 cm with in width, 50 cm long and placed at a height of 55 cm. At the beginning of each test, the animals were allowed to get used to the experimental room for 30 minutes and, at the beginning of the procedure they were placed in the center of the apparatus in front of an open arm. The animals were allowed to explore the apparatus for 5 min. The video signal was digitized and analyzed with the EthoVision XT 8.5 software (Noldus Information Technology, Leesburg, VA). An open arm entry was considered when the center of the mouse is in one of the open arms. The device was cleaned with ethanol before and between animals.

**Congo-red binding**
For the spectroscopic test with CR, a working solution at 40 µM in buffer (5 mM potassium phosphate, 150 mM NaCl, pH 7.4) was prepared and filtered immediately before use. 5 µL of protein (10 µM, incubated until the corresponding time in each case) was added to 5 µL of the stock CR solution, so that a final ratio of 1:2

(protein:CR) is maintained. The UV-visible spectrum was recorded in a spectrophotometer (BioSpectrometer, Eppendorf) between 400 and 700 nm, collecting the absorbance values at 480 and 540 nm to subsequently calculate the concentration of bound CR [3,4].

#### *References for supplementary material*


1. Crawley, J. N. Behavioral phenotyping of transgenic and knockout mice: Experimental design and evaluation of general health, sensory functions, motor abilities, and specific behavioral tests. *Brain Res.* **835**, 18–26 (1999).

2. Kayed, R. *et al.* Fibril specific, conformation dependent antibodies recognize a generic epitope common to amyloid fibrils and fibrillar oligomers that is absent in prefibrillar oligomers. *Mol. Neurodegener.* **2**, 1–11 (2007).

3. Wurth, C., Guimard, N. K. & Hecht, M. H. Mutations that reduce aggregation of the alzheimer°Fs Aβ42 peptide: An unbiased search for the sequence determinants of Aβ amyloidogenesis. *J. Mol. Biol.* **319**, 1279–1290 (2002).

4. Hervás, R. *et al.* Common features at the start of the neurodegeneration cascade. *PLoS Biol.* **10**, e1001335 (2012).
